# Inverted translational control of eukaryotic gene expression by ribosome collisions

**DOI:** 10.1101/562371

**Authors:** Heungwon Park, Arvind R. Subramaniam

## Abstract

The canonical model of eukaryotic translation posits that efficient translation initiation increases protein expression and mRNA stability. Contrary to this dogma, we show that increasing initiation rate can decrease both protein expression and stability of certain mRNAs in the budding yeast, *S. cerevisiae*. These mRNAs contain a stretch of poly-basic residues that cause ribosome stalling. Using computational modeling, we predict that the observed decrease in gene expression at high initiation rates occurs when ribosome collisions at stalls stimulate abortive termination of the leading ribosome and cause endonucleolytic mRNA cleavage. We test our prediction by identifying critical roles for the collision-associated quality control factors, Asc1 and Hel2 (RACK1 and ZNF598 in humans, respectively). Remarkably, hundreds of *S. cerevisiae* mRNAs that contain ribosome-stall sequences also exhibit lower translation efficiency. We propose that these mRNAs have undergone evolutionary selection for inefficient initiation to escape collision-stimulated reduction in gene expression.

## Introduction

Translation of mRNAs by ribosomes is a critical regulatory point for controlling eukaryotic gene expression. Initiation is usually the slowest kinetic step in the translation of eukaryotic mRNAs ^1^. Therefore, increasing the initiation rate of mRNAs typically results in higher expression of the encoded protein. Higher initiation rates can also protect eukaryotic mRNAs from decay. This occurs primarily through preferential binding of translation initiation factors over mRNA decay factors to the 5′ cap of mRNAs ^2–4^. Such mRNA stabilization amplifies the positive effect of high initiation rate on protein expression ^5, 6^. Thus, efficient initiation is widely associated with increased mRNA stability and higher protein expression ^7–9^.

Efficient initiation is also required for quality control at ribosomes that slow down or stall during elongation ^10, 11^. Slowing down of ribosomes at non-optimal codons accelerates mRNA decay ^12–14^. The No-go mRNA decay (NGD) and the ribosome-associated quality control (RQC) pathways target ribosomes stalled at poly-basic stretches, rare codon repeats, or mRNA stem loops ^15, 16^. These pathways together mediate cleavage of mRNAs, degradation of nascent peptides, and recycling of ribosomes and peptidyl-tRNAs ^17–23^. Mutations that destabilize mRNAs or block translation initiation also suppress the above quality control events ^11, 15, 24^. This raises the question of how normal translation and quality control compete with each other to set the overall stability and protein expression of stall-containing mRNAs.

Computational models of translation have been extensively used to investigate the effect of ribosome kinetics on protein expression ^25, 26^. The widely used traffic jam model predicts higher protein expression as initiation rate is increased until elongation becomes limiting ^27, 28^. In the elongation-limited regime of the traffic jam model, queues of collided ribosomes prevent further increase in protein expression ^29^. However, this elongation-limited regime of queued ribosomes or the associated constancy in protein expression has not been observed in experiments. On the contrary, recent experiments reveal that collided ribosomes trigger quality control processes such as mRNA cleavage and degradation of the nascent peptide ^24, 30, 31^. Computational models that accurately capture these kinetic events at stalled ribosomes might reveal new modes of regulating eukaryotic gene expression.

In this study, we systematically characterized the effect of initiation rate on protein expression and stability of both normal and stall-containing mRNAs in *S. cerevisiae*. By designing 5′ UTR mutations that de-couple translation initiation rate from normal mRNA stability, we uncover that high initiation rates can reduce gene expression of stall-containing mRNAs. To our knowledge, this inverse relation has not been previously observed in experiments. Our theory predicts that high initiation rates decrease protein expression and mRNA stability when ribosome collisions cause the leading ribosome to abort translation or cleave the mRNA. We tested our prediction by measuring the effect of various known quality control factors on protein expression and mRNA stability. Strikingly, Hel2 and Asc1, the only two identified factors associated with ribosome collisions ^24, 30, 31^, are also required for the decrease in gene expression at high initiation rates. We show that hundreds of endogenous *S. cerevisiae* mRNAs with ribosome stalls have unusually low translation efficiency. We propose that these mRNAs are evolutionarily selected for inefficient initiation to escape from collision-stimulated reduction in gene expression.

## Results

### 1. High initiation rates decrease protein output of stall-containing *S. cerevisiae* mRNAs.

To measure the effects of initiation rate variation on protein expression, we used a fluorescent reporter system in the budding yeast, *S. cerevisiae* (Fig. 1A). Our reporters consist of the *PGK1* gene from *S. cerevisiae* that has been extensively used in previous studies of mRNA translation and stability ^5, 13^. We fused the *PGK1* coding sequence to a 3×FLAG-encoding sequence at the 5′ end and the yellow fluorescent protein (*YFP*) gene at the 3′ end. We integrated this 2kb protein-coding region, under the control of the *GPD* promoter, the *GPD* 5′ UTR and the *CYC1* 3′ UTR, as a single copy into the genome (Methods). We expressed a red fluorescent protein (mKate2) gene with identical regulatory sequences as our reporter from a different genomic locus for normalizing the measured YFP fluorescence.

**Figure 1.**
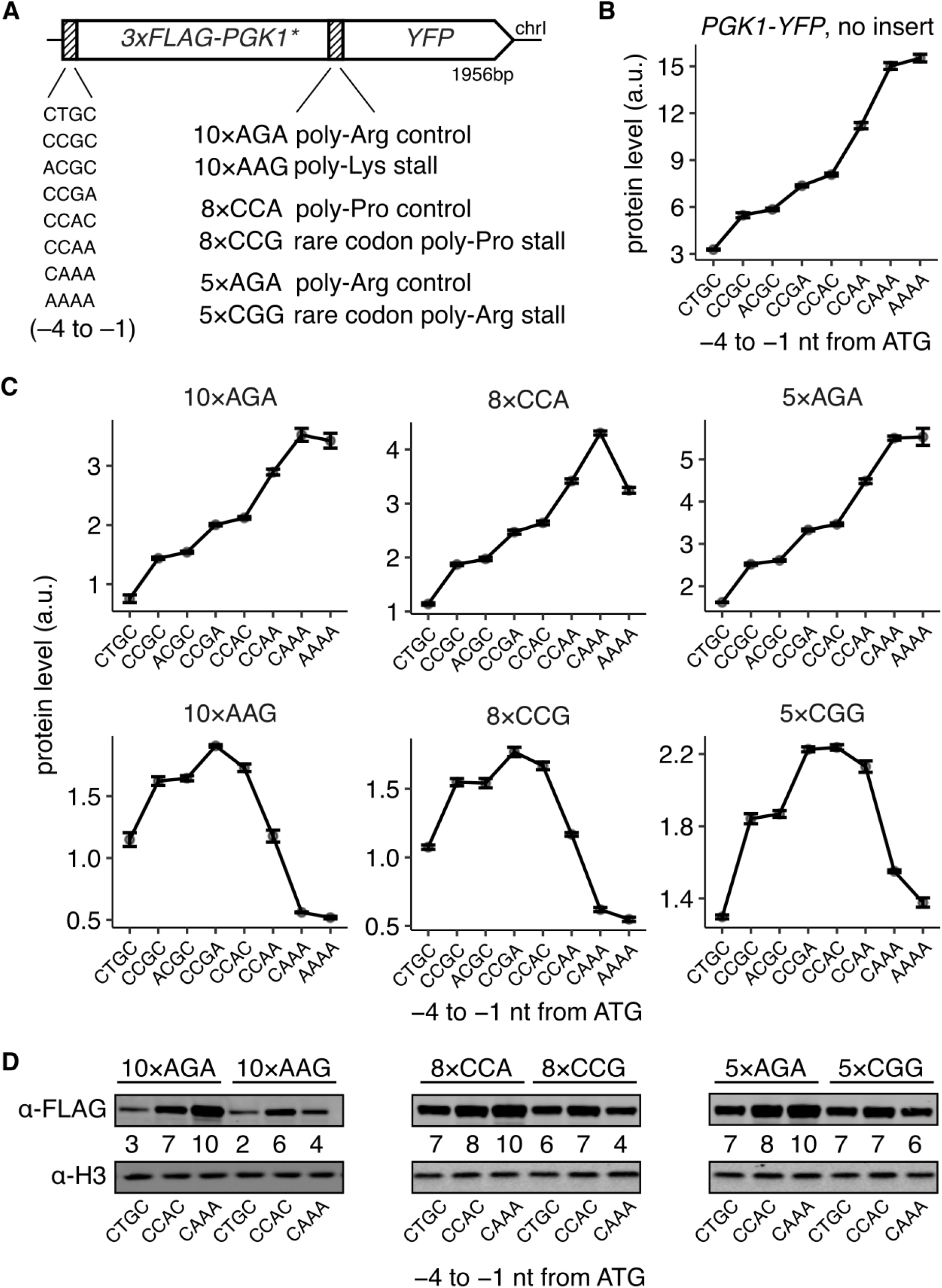
High initiation rates decrease protein output of stall-containing *S. cerevisiae* mRNAs. **(A)** Schematic of *3×FLAG-PGK1*-YFP* reporters used in C and D. The −4 to −1 nt region preceding the ATG start codon had one of 8 different Kozak sequences as indicated. One of three different stall sequences or their respective controls were inserted at the end of *PGK1\**. *PGK1\** was modified from wild-type *PGK1* sequence based on a previous study ^11^ (see Methods). **(B)** Protein levels of *3×FLAG-PGK1-YFP* control reporters with no insert and wild-type *PGK1* sequence. **(C)** Protein levels of the constructs shown in A. (*Caption continued on next page.*) Protein levels are quantified as the mean YFP fluorescence of 10,000 cells for each strain as measured using flow cytometry. Protein levels are expressed as arbitrary units (a.u.) relative to the mean *RFP* levels from a constitutively expressed mKate2 control. Error bars show standard error of the mean over 3 or 4 independent transformants in B and C. Many error bars are smaller than data markers. 2 of the total 192 strains were clear outliers and removed after manual inspection. **(D)** Western blots of reporters with low (CTGC), medium (CCAC), or high (CAAA) initiation rates and with indicated stall sequences or controls using antibody against the FLAG epitope at the N-terminus. Histone H3 level is shown as loading control. Numbers for each lane indicate the ratio of the FLAG signal to the H3 signal, and are normalized to a maximum of 10 within each blot.

To confirm our reporter system’s utility for studying both normal and stall-containing mRNAs, we introduced 5 tandem arginine AGA codons (5×AGA) at five different locations along the *PGK1* gene (henceforth referred to as *PGK1\**, Fig. S1A), following earlier work ^11^. We individually mutated each of the five 5×AGA repeats to their synonymous CGG rare arginine codons and measured their effect on YFP expression using flow cytometry. The 5×CGG rare codons reduce YFP expression by 40–70%, with the stalls away from the ATG start codon having a stronger effect (Fig. S1B).

To vary the initiation rate of our reporters, we designed 8 different 5′ UTR variants where we mutate the 4 nt preceding the ATG start codon. These mutations vary 3×FLAG-PGK1-YFP expression over a 5-fold range (Fig. 1B). This hierarchy of YFP expression among the 5′ UTR variants is concordant with previous measurements ^32^ (Fig. S1C). Since the 5′ UTR mutations are located over 30 nt away from the 5′ end of the mRNA, we expect them to have minimal effects on the binding of translation initiation factors and mRNA decay factors to the 5′ cap.

We then combined the 5′ UTR mutations with various stall and control sequences to measure their combinatorial effect on protein expression (Fig. 1A). We used the 5×CGG stall sequence from above, as well as 8×CCG rare proline codon repeats and 10×AAG lysine codon repeats that are known to trigger ribosome stalling ^33–35^. As controls, we used 5×AGA, 8×CCA, and 10×AGA repeats, respectively.

Strikingly, the three 5′ UTR variants (CCAA, CAAA, AAAA) that have up to 2-fold higher protein level than the median for normal mRNAs (Fig. 1B), instead cause up to a 4-fold reduction in protein level when the mRNA contains any of the three ribosome stalls (lower three panels in Fig. 1C). This decrease in protein expression at high initiation rates is largely absent when the stalls were replaced by control sequences of synonymous common codons (5×AGA and 8×CCA, Fig. 1C). Similarly, replacing the 10×AAG lysine repeat stall by a 10×AGA arginine control abrogates the decrease in protein expression at high initiation rates (Fig. 1C, left panels). Importantly, the 10×AAG repeat differs from the 10×AGA control only by a single nucleotide insertion and deletion, and is located over 1.3 kb away from the initiation codon. Hence, changes in mRNA secondary structure or other subtle physical interactions between the 5′ UTR mutations and the stall mutations are unlikely to account for the dramatically different behaviors of the 10×AAG and 10×AGA inserts at high initiation rates. Further, since AAG is a common lysine codon with abundant cellular tRNA ^36, 37^ and our reporter is integrated as a single copy in the *S. cerevisiae* genome, the observed effects on protein expression at high initiation rates are unlikely to be caused by titration of rare tRNAs by the stall-encoding repeats. Finally, western blotting against the N-terminus 3×FLAG epitope qualitatively reproduces the differences in protein expression as measured by flow cytometry (Fig. 1D). These results confirm that high initiation rates have qualitatively distinct effects on expression of the full length protein from reporter mRNAs with or without ribosome-stall sequences.

### 2. High initiation rates decrease lifetime of stallcontaining *S. cerevisiae* mRNAs.

We reasoned that the change in protein expression with initiation rate in our experiments could arise from change in mRNA stability, change in the rate at which ribosomes finish synthesis of the full length protein from each mRNA, or a combination of both. To isolate the effect of initiation rate variation on mRNA stability, we developed a high throughput sequencing-based assay to quantify mRNA levels of our *PGK1-YFP* reporters at high resolution (Fig. 2A). In our as-say, each combination of 5′ UTR mutation and codon repeat is tagged with four distinct 8 nt barcodes in their 3′ UTR. We pooled these barcoded reporters and integrated them into the genome of *S. cerevisiae*. Reporter mRNA from the pooled strains was reverse-transcribed to cDNA. The barcode region from both cDNA and genomic DNA was then amplified by PCR and counted by sequencing. The relative mRNA level of each reporter variant was obtained by normalizing the cDNA count of the corresponding barcode by its genomic DNA count.

**Figure 2.**
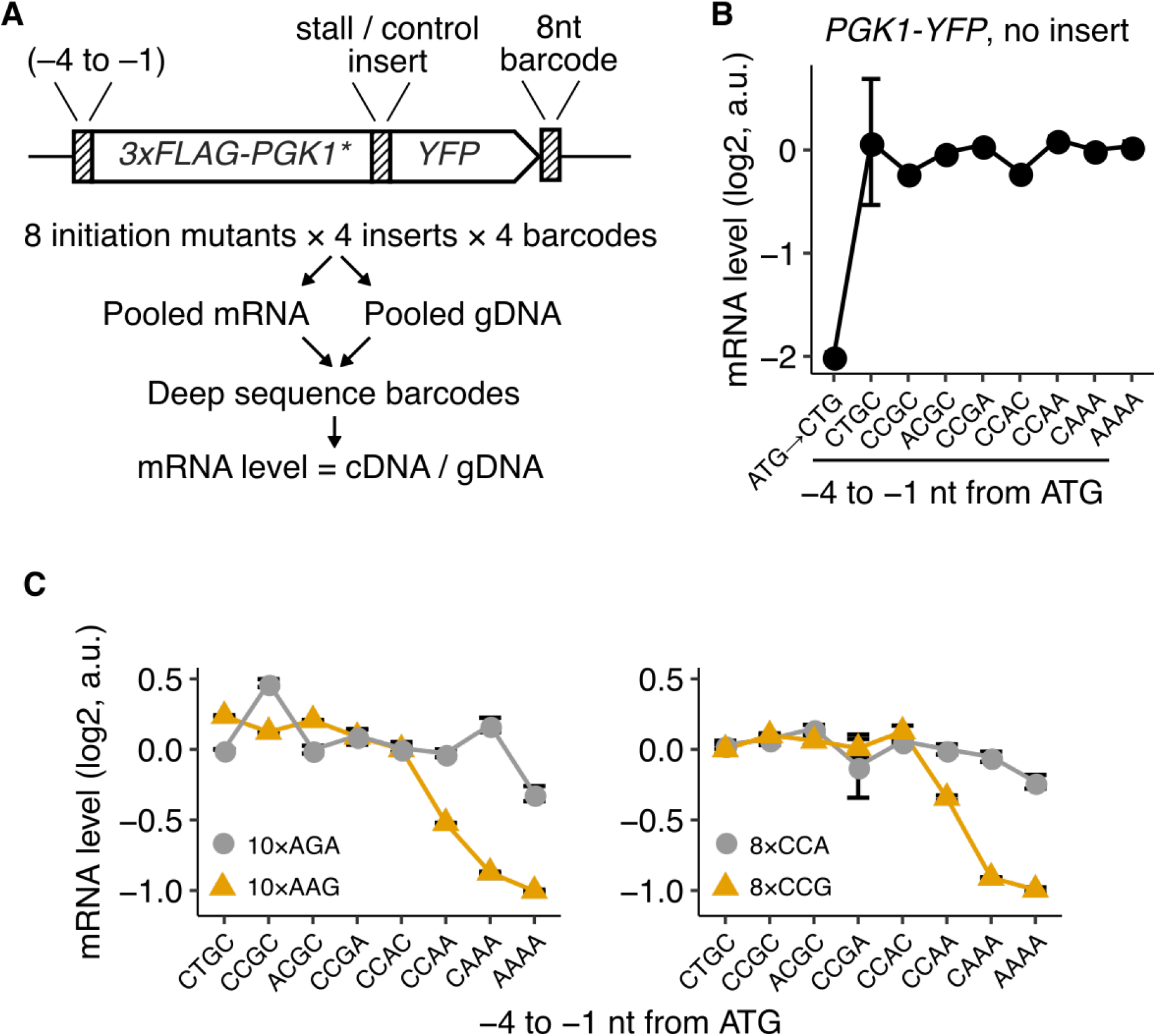
High initiation rates decrease stability of stall-containing *S. cerevisiae* mRNAs. **(A)** Schematic of deep-sequencing assay used for measuring mRNA levels. Reporters with different 5′ UTR mutations and stall / control inserts were tagged with four unique 8 nt barcodes in their 3′ UTR. The reporters were pooled and transformed into *S. cerevisiae*. cDNA and genomic DNA (gDNA) from the pooled reporters were amplified and the barcodes counted by high-throughput sequencing. mRNA level was calculated as cDNA counts normalized by gDNA counts for each barcode, and median-normalized within each set of reporters with varying 5′ UTR mutations. **(B)** mRNA levels of *3×FLAG-PGK1-YFP* control reporters with no insert and wild-type *PGK1* sequence. The left-most point is for a reporter with the ATG start codon mutated to CTG. **(C)** mRNA levels of *3×FLAG-PGK1*-YFP* constructs with stall / control inserts. Error bars in (B) and (C) show standard error over 3 or 4 distinct barcodes for each reporter variant. Most error bars are smaller than data markers.

We validated our experimental strategy by measuring steady-state mRNA levels of the *3×FLAG-PGK1-YFP* reporters with no ribosome stall sequences and eight different 5′ UTR mutations. mRNA levels differ less than 25% between the eight 5′ UTR variants (Fig. 2B), even though protein levels differ more than 5-fold between these variants (Fig. 1B). Since steady state mRNA levels reflect the balance between mRNA synthesis and decay, we parsimoniously interpret these measurements to imply that our 5′ UTR mutations change only the translation initiation rate at the ATG start codon of our reporters without affecting their mRNA stability or transcription rate. As a positive control, mutating the ATG start codon of our reporter to CTG decreases mRNA levels by 4-fold (Fig. 2B). We expect this decrease if absence of the main ATG start codon results in translation initiation at one of the four downstream out-of-frame start codons in the 3×FLAG-encoding region, which then destabilizes the mRNA by NMD.

We then applied our experimental strategy to measure mRNA levels of the 5′ UTR variants of our *3×FLAG-PGK1*-YFP* reporters with various stall or control inserts (Fig. 2C). The 8×CCG and 10×AAG stall sequences result in up to a 2-fold decrease in mRNA levels specifically in the high initiation rate regime (yellow triangles, Fig. 2C). By contrast, *3×FLAG-PGK1*-YFP* reporters with the 8×CCA and 10×AGA control inserts show little or no decrease in mRNA levels at high initiation rates (grey circles, Fig. 2C), which is similar to the *3×FLAG-PGK1-YFP* constructs without inserts (Fig. 2B). We interpret the decrease in mRNA levels of the 8×CCG and 10×AAG stall-containing reporters at high initiation rates as changes in their mRNA lifetime. This interpretation is justified given that the 5′ UTR mutations, on their own, do not affect mRNA levels (Fig. 2B). We were unable to directly confirm this interpretation because established approaches for measuring mRNA lifetimes rely on changes in growth medium or temperature^4, 11^. We found that these perturbations affect cell growth rate, which presumably alters the global translation initiation rate and thus ablates the reporter-specific effect of high initiation rates (data not shown).

Comparison between the protein and mRNA measurements from our *PGK1-YFP* reporters show that protein levels at high initiation rate decrease up to 4-fold from their peak value, while mRNA levels decrease only 2-fold (CCGA vs. AAAA 5′ UTR mutations in Fig. 1C and 2C). Thus, we conclude that the decrease in protein expression at high initiation rates arises from both steady-state change in mRNA levels, as well as change in the synthesis rate of the full length protein from each mRNA.

### 3. Collision-stimulated abortive termination model predicts reduced protein output at high initiation rates

We hypothesized that computational modeling can provide mechanistic insight into how high initiation can result in lower protein expression and reduced stability of stall-containing eukaryotic mRNAs. Since kinetic models that consider both translation and stability of eukaryotic mRNAs have not been formulated so far to our knowledge, we first defined a joint model for normal translation and canonical mRNA decay in eukaryotes (Methods). We modeled normal translation as a three step process composed of initiation, elongation, and termination, similar to previous studies ^38, 39^. Each ribosome occupies a footprint of ten codons on mRNAs ^40^, and initiation and elongation occur only if the required footprint is not blocked by another leading ribosome. We modeled canonical mRNA decay in eukaryotes as a three step process of deadenylation, decapping, and exonucleolysis ^41^. Even though a more detailed kinetic model of canonical mRNA decay is available ^41^, since we aimed to predict overall mRNA lifetime, we considered only the three main steps of canonical mRNA decay. We chose the rate parameters for these decay steps ^41^ (Table S3) to result in a mean mRNA lifetime of 35 minutes corresponding to that of the *PGK1* mRNA in *S. cerevisiae* ^5^. We used a rule-based modeling approach and performed exact stochastic simulations using an agent-based framework to predict mRNA lifetime and protein expression (Methods).

We then systematically investigated the effect of quality control at elongation stalls on protein expression. We first studied kinetic models of how stalled ribosomes undergo abortive termination by setting the mRNA decay rate to zero (Table S3). In our model, abortive termination includes all molecular processes that terminate polypeptide synthesis including splitting of the stalled ribosome, degradation of the nascent peptide, and hydrolysis of the peptidyl-tRNA. We did not distinguish between these different molecular events since all of them reduce synthesis rate of the full length protein, which is the quantity that we measure in our experiments.

We considered three kinetic models of how abortive termination might occur at stalled ribosomes from intact mRNAs (Fig. 3A, Methods). In the simple abortive termination (SAT) model, stalled ribosomes are subject to kinetic competition between abortive termination and normal elongation (Fig. 3A). This SAT model has been considered in recent modeling studies ^42–44^. Since recent experiments found a role for ribosome collisions in quality control ^24, 30, 31^, we then considered two possible effects of ribosome collisions on abortive termination. In the collision-stimulated abortive termination (CSAT) model, only stalled ribosomes that get hit by trailing ribosomes undergo abortive termination (Fig. 3A). Conversely, in the collide and abortive termination (CAT) model, only trailing ribosomes that run into leading stalled ribosomes undergo abortive termination (Fig. 3A). Finally, as a control, we also considered the traffic jam (TJ) model in which ribosomes do not undergo abortive termination (Fig. 3A).

**Figure 3.**
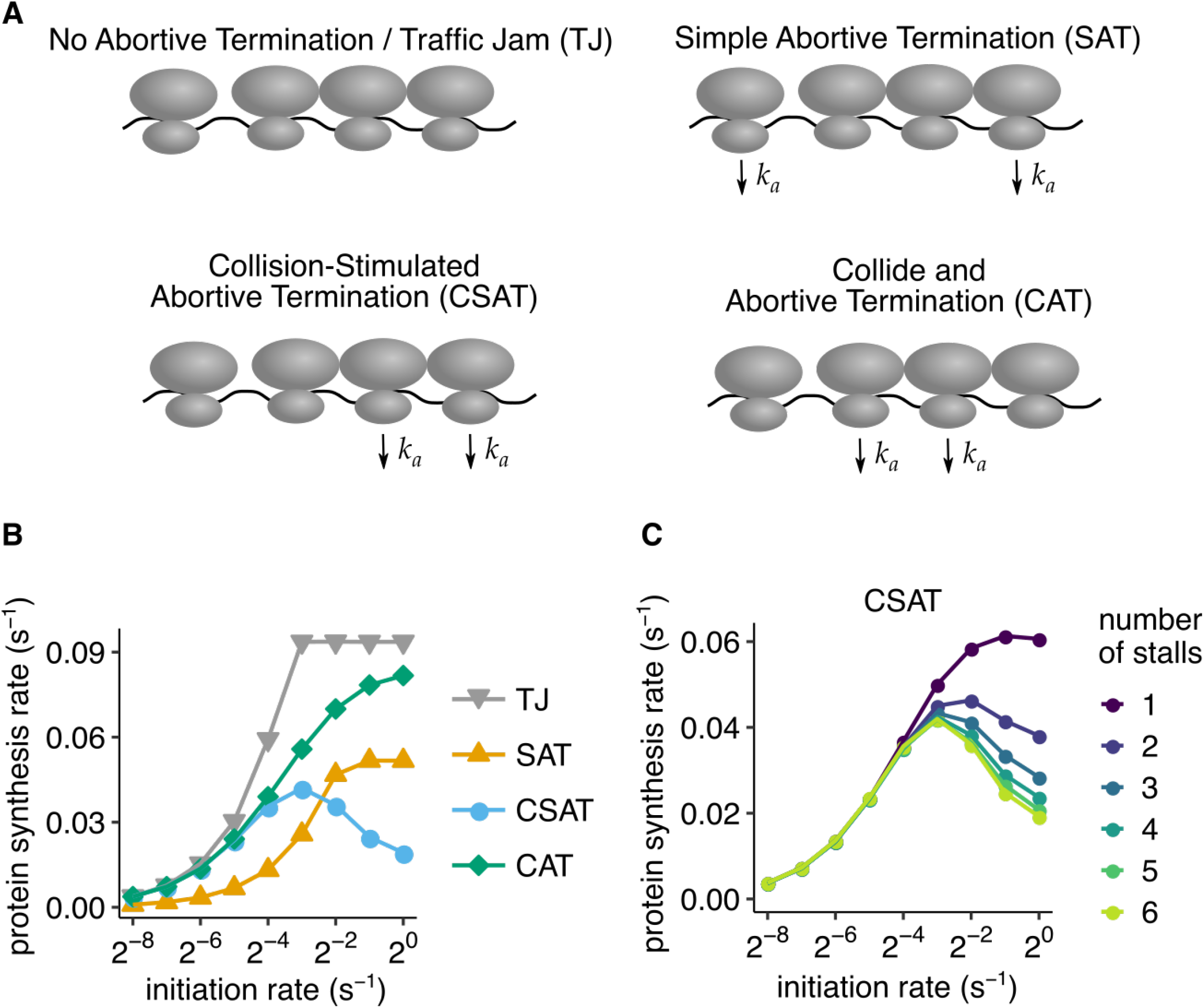
Collision-stimulated abortive termination model predicts reduced protein output at high initiation rates. **(A)** Schematic of different abortive termination models simulated in (B). *k*_*a*_ denotes the non-zero rate of abortive termination of ribosomes from indicated configurations. In the TJ model, ribosomes do not abort translation. In the SAT model, only ribosomes that have not undergone collision from the mRNA entry side abort. In the CAT model, only ribosomes that have undergone collision from the mRNA entry side (‘trailing’ ribosomes in a collision) abort. In the CSAT model, only ribosomes that have undergone collision from the mRNA exit side (‘leading’ ribosomes in a collision) abort. **(B)** Average protein synthesis rate as a function of initiation rate predicted using stochastic simulations of the four models in (A). The simulations were of a 650 codon mRNA corresponding to the *3×FLAG-PGK1*-YFP* reporters in our experiments. Ribosome stalls were simulated as a consecutive stretch of 6 slowly translated codons with a net elongation rate of 0.1*s*^−1^. All other codons had an elongation rate of 10*s*^−1^. The stalls were located after 400 codons, corresponding to their approximate location in our experiments. **(C)** Effect of varying the number of tandem (consecutive) slowly-translated codons encoding the stall in the CSAT model. The elongation rate of each slowly translated codon was scaled to maintain a net elongation rate of 0.1*s*^−1^ across the stall. mRNA decay rate was set to zero in these simulations to isolate the effect of abortive termination. All other model parameters are listed in Table S3. Standard error from repeating stochastic simulations with different initial random seeds are smaller than data markers in (B) and (C).

We simulated the above described kinetic models of abortive termination with varying initiation rate and analyzed the predicted protein synthesis rate (defined as the number of full proteins produced per second). We simulated elongation stalls by introducing a stretch of 6 tandem poorly translated codons after 400 codons in a 650 codon mRNA, which is similar in length to the *PGK1-YFP* reporters that we use in our experiments. In our simulations, the control traffic jam (TJ) model without abortive termination exhibits a linear increase in protein synthesis rate with initiation rate until the elongation rate at the stall becomes rate-limiting (grey triangles, Fig. 3B; S2A). This observation is consistent with previous studies of this model ^45, 46^. The simple abortive termination (SAT) model exhibits a similar behavior to the TJ model, but with lower protein synthesis rate and a higher initiation rate at which protein synthesis rate reaches saturation (yellow triangles, Fig. 3B, S2B, S2C).

Unexpectedly, the two kinetic models in which ribosome collisions cause abortive termination of either the trailing or the leading ribosome (Fig. 3A) exhibit different behaviors as initiation rate is varied. In the CAT model in which the trailing ribosome abortively terminates upon collision, protein synthesis rate increases monotonically as the initiation rate is increased (green diamonds, Fig. 3B). By contrast, in the CSAT model in which the leading ribosome abortively terminates upon collision, protein synthesis rate increases initially, then reaches a maximum, and surprisingly decreases with further increase in initiation rate (blue circles, Fig. 3B).

The above dichotomy between the CSAT and CAT models of abortive termination at ribosome collisions can be intuitively understood as follows: In the CAT model, even though the trailing ribosomes at collisions undergo abortive termination, every ribosome that reaches the stall eventually finishes protein synthesis at a rate determined by the elongation rate at the stall. In the CSAT model, the leading stalled ribosome is stimulated to undergo abortive termination upon collision; thus, at sufficiently high initiation rates and abortive termination rates, very few ribosomes will get past the stall. Stated differently, the difference between the CSAT and CAT models arises because all ribosomes that finish protein synthesis stall at the slow codons, while not all ribosomes collide with a leading ribosome before they finish protein synthesis.

The decrease in protein expression at high initiation rates in the CSAT model is counter-intuitive, and to further probe its origin, we systematically varied the elongation rate at the stall and the number of tandem (consecutive) codons encoding the stall. The decrease in protein synthesis rate requires the initiation rate to exceed the total elongation rate past the stall (Fig. S2D) since ribosome collisions occur only in this regime. More surprisingly, the decrease in protein synthesis occurs only if ribosomes stall at multiple consecutive codons (Fig. 3C). As a concrete example, the CSAT model predicts that 2 tandem stall codons with 5 *s* ribosome dwell time at each codon decreases protein synthesis rate at high initiation rates, but a single stall codon with 10 *s* ribosome dwell time does not. This non-intuitive prediction can be understood by observing that the number of kinetic partitions between normal elongation and abortive termination changes dynamically with initiation rate in the CSAT model with *n* consecutive stall codons. At low initiation rates (*k*_*init*_ ≪ *k*_*stall*_), there is no kinetic partitioning towards abortive termination, similar to the TJ model. At high initiation rates (*k*_*init*_ ≫ *k* _*stall*_), each ribosome that finishes protein synthesis undergoes *n* kinetic partitions towards normal elongation, similar to the SAT model. However, at intermediate initiation rates (*k*_*init*_ ∼ *k*_*stall*_), the average number of kinetic partitions increases from 0 to *n* as the initiation rate is increased. In this sense, the CSAT model ‘interpolates’ between the TJ model and the SAT model.

Thus, our simulations of different kinetic models of abortive termination reveal distinct signatures of initiation rate variation on protein expression. The CSAT model, unlike the other models, predicts a decrease in protein expression at high initiation rate that matches the observation from our experiments (Fig. 1C). Importantly, our simulations reveal that this decrease is not simply a consequence of ribosome collisions stimulating quality control, but it has two other essential ingredients (Fig. 3B, C): The leading ribosome in the collision undergoes abortive termination, and the stall itself is composed of multiple kinetic steps.

### 4. Collision-stimulated endonucleolytic cleavage model predicts decrease in mRNA lifetime at high initiation rates

Since endonucleolytic cleavage has been shown to reduce mRNA lifetime of stall-containing mRNAs ^15^, we next considered how different kinetic models of how endonucleolytic mRNA cleavage at ribosome stalls decreases mRNA lifetime as a function of initiation rate (Fig. 4A, Methods). We modeled endonucleolytic mRNA cleavage as occurring on the 5′ side of the stalled ribosome ^47, 48^. In our modeling, once mRNAs are endonucleolytically cleaved, translation initiation is immediately repressed through decapping, and ribosomes that have already initiated on the 5′ mRNA fragment are efficiently recycled once they reach the truncated 3′ end ^19, 20^. We simulated ribosome stalling by introducing a stretch of 6 poorly translated codons after 400 codons in a 650 codon mRNA, which is similar to our *PGK1-YFP* reporters. To isolate the effect of endonucleolytic cleavage on our predictions, we set the abortive termination rate from intact mRNAs to be zero. We monitored the mean lifetime of mRNAs, which we define as the time interval between the end of transcription and the start of 5′–3′ exonucleolytic decay after decapping. As before, we also monitored protein synthesis rate as we varied the initiation rate of mRNAs in our simulations.

**Figure 4.**
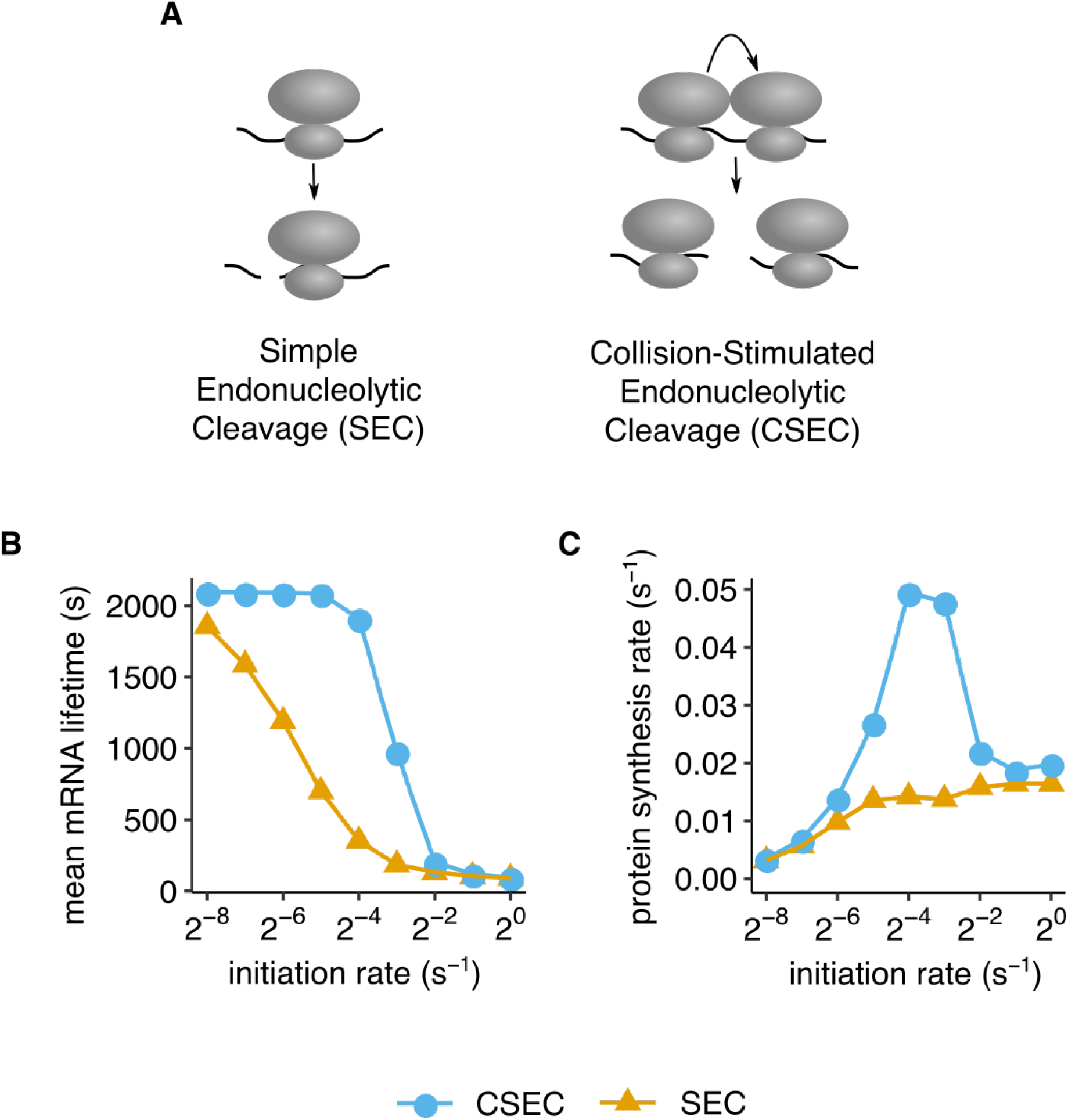
Collision-stimulated endonucleolytic cleavage model predicts decrease in mRNA lifetime at high initiation rates. **(A)** Schematic of simple collision-independent (SEC) and collision-stimulated (CSEC) endonucleolytic mRNA cleavage models simulated in (B) and (C). **(B, C)** Average mRNA lifetime and protein synthesis rate over 10^6^*s* as a function of initiation rate predicted using stochastic simulations of the models in (A). Reporters were simulated as in Fig. 3, but with a non-zero endonucleolytic cleavage rate of 0.001*s*^−1^ and a net elongation rate of 0.1*s*^−1^ across the stall. Canonical mRNA decay was allowed while abortive termination rate was set to zero. All other model parameters are listed in Table S3. Standard error from repeating stochastic simulations with different initial random seeds are smaller than data markers in (B) and (C).

We considered two distinct kinetic models of how endonucleolytic mRNA cleavage might occur at ribosome stalls (Fig. 4A, Methods). In the simple endonucleolytic cleavage model (SEC), cleavage occurs through simple kinetic competition with normal elongation ^49^. In the collision-stimulated endonucleolytic cleavage model (CSEC), cleavage occurs only between two collided ribosomes.

Both the simple and collision-stimulated models of endonucleolytic cleavage predict a decrease in mRNA lifetime at high initiation rates (Fig. 4B). However, while the SEC model predicts a gradual decrease in mRNA lifetime (yellow triangles, Fig. 4B), the CSEC model predicts a sharp decrease in mRNA lifetime as the initiation rate is increased (blue circles, Fig. 4B). The dependence of mRNA lifetime on initiation rate in the SEC model is surprising since there is no direct role for initiation rate in this model. However, even in the SEC model, the endonucleolytic cleavage probability of a given mRNA increases with the frequency of ribosome stalling, which in turn increases with the frequency of initiation. Thus in both our kinetic models of endonucleolytic cleavage, mRNAs transition from being primarily degraded through the canonical decay pathway at low initiation rates to being endonucleolytically cleaved at high initiation rates.

Even though the mRNA lifetime decreases at high initiation rate in both the simple and collision-stimulated cleavage models, the protein synthesis rate in these models display strikingly different behaviors as a function of initiation rate (Fig. 4C). In the SEC model, protein synthesis rate increases monotonically with initiation rate (yellow triangles, Fig. 4C): At low initiation rates, it increases linearly where mRNAs are degraded primarily through the canonical decay pathway, and saturates at high initiation rates at a value determined by the endonucleolytic cleavage rate at stalls (Fig. S3A). In the high initiation rate regime, each mRNA is translated by a fixed number of ribosomes (on average) before it undergoes endonucleolytic cleavage in the SEC model. By contrast, in the CSEC model, protein synthesis rate exhibits a non-monotonic behavior (blue circles, Fig. 4C): It increases linearly until the initiation rate matches the elongation rate at the stall, at which point it decreases sharply and saturates at the same value as the SEC model. Unlike the CSAT model (blue circles, Fig. 3C), the behavior of the CSEC model does not depend sensitively on the number of stalling codons that the ribosome transits through (Fig. S3B).

Thus, our simulations of different kinetic models of endonucleolytic mRNA cleavage reveal distinct signatures of initiation rate variation on mRNA lifetime and protein expression. The CSEC model, unlike the SEC model, predicts the decrease in mRNA stability observed in our experiments only at high initiation rates (Fig. 2C). The CSEC model also correctly predicts the observed decrease in protein expression at high initiation rates (Fig. 1C), which is independent of the effect of abortive termination on protein expression.

### 5. Hel2 and Asc1 attenuate translation of stall-containing mRNAs only at high initiation rates

We then sought to test the prediction from our modeling that collision-stimulated quality control (CSEC and CSAT) directly contributes to the decrease in protein expression and mRNA stability at high initiation rates from stall-containing mRNAs. Towards this, we focused on several factors that have been previously implicated in quality control at ribosome stalls, but whose effect on gene expression as a function of initiation rate has not been characterized. The ribosome-ubiquitin E3 ligase Hel2 (ZNF598 in humans) is activated by collided ribosomes ^24, 30, 31^. The ribosome-associated protein Asc1 (RACK1 in humans) is located at the interface between collided ribosomes ^30, 31^, and along with Hel2, couples translational arrest to nascent chain degradation ^50–53^. We also consider the ribosome recycling factor Dom34 (PELO in humans) that plays a critical role in No-go decay ^15, 49, 54^. Nascent chain degradation is mediated by the E3 ligase Ltn1 (LTN1 in humans), which is part of the RQC complex ^16, 18, 55^. We measured protein expression and mRNA levels from our *3×FLAG-PGK1*-YFP* reporters in *S. cerevisiae* strains in which *HEL2, ASC1, DOM34* and *LTN1* were individually deleted (Fig. 5A–C).

**Figure 5.**
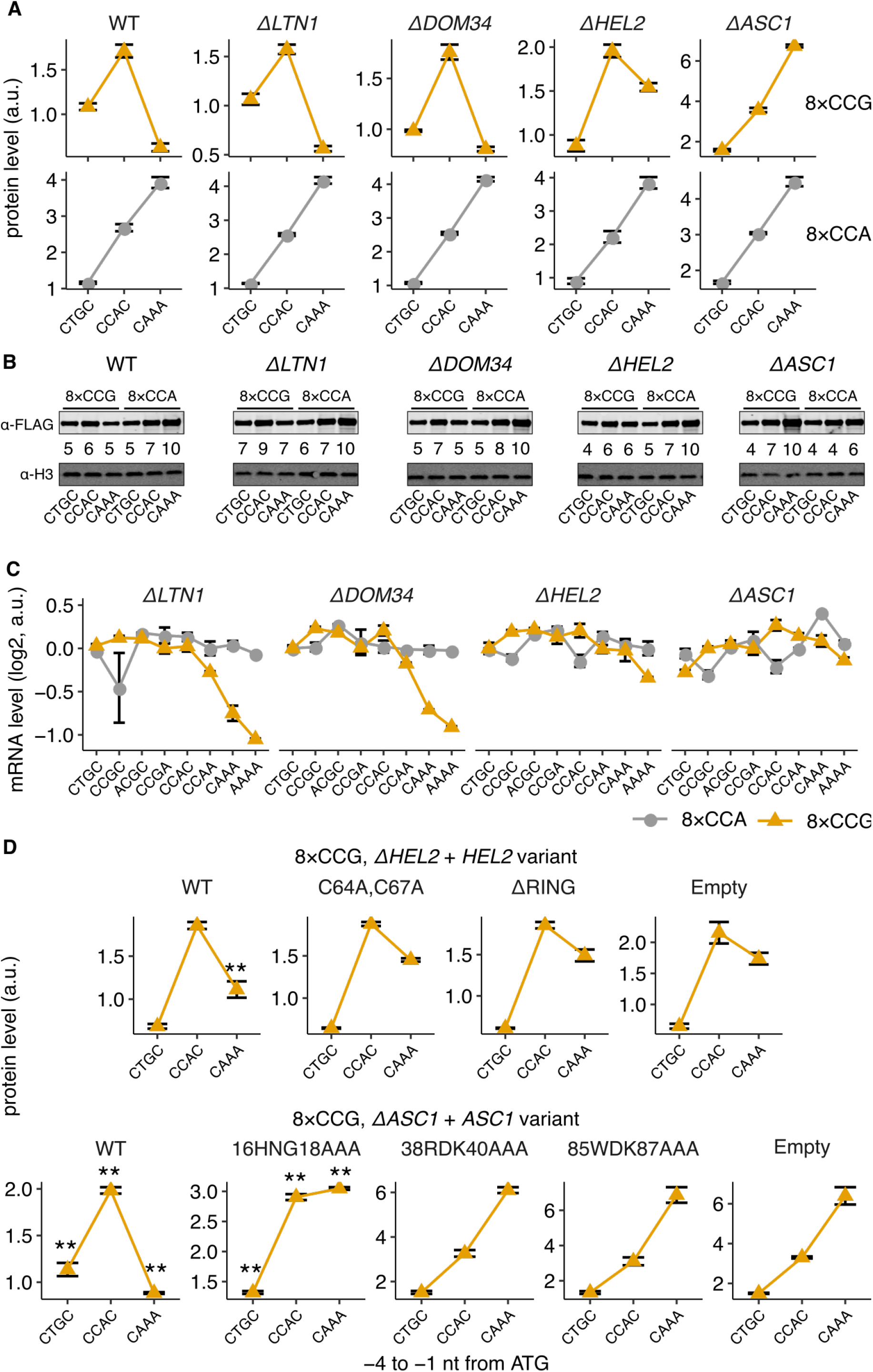
Hel2 and Asc1 attenuate translation of stall-containing mRNAs only at high initiation rates. (*Caption continued on next page.*) **(A)** Protein levels of *3×FLAG-PGK1*-YFP* reporters (see Fig. 1A) with low (CTGC), medium (CCAC), or high (CAAA) initiation rates and with stall (8×CCG) or control (8×CCA) repeats. The reporters were integrated into the genome of either the wild-type strain (WT), or isogenic strains with full deletions of *LTN1, DOM34, HEL2*, or *ASC1*. **(B)** Western blots of reporters from (A) using antibody against the FLAG epitope at the N-terminus. Histone H3 level is shown as loading control. Numbers for each lane indicate the ratio of the FLAG signal to the H3 signal, and normalized to a maximum of 10 within each blot. **(C)** mRNA levels of the *3×FLAG-PGK1*-YFP* reporters with the indicated 5′ UTR mutations and stall sequences. mRNA levels were quantified as in ig. 2A. Error bars show standard error over 3 or 4 distinct barcodes for each reporter variant. **(D)** Protein levels of the 8×CCG stall reporter expressed in either *δHEL2* (top) or *δASC1* (bottom) strain and complemented with the indicated Hel2 or Asc1 variant, respectively. Error bars in (A) and (D) show standard error over 3 or 4 independent transformants. Many error bars are smaller than data markers. ** in (D) denotes P < 0.01 (two-tailed *t*-test) for comparison between indicated Hel2/Asc1 variant and ‘Empty’ for each of the three –4 to –1 5′ UTR sequences. Absence of ** indicates P > 0.05.

Deletion of *ASC1* completely rescues protein expression from the 8×CCG stall-containing reporters (*ΔASC1* in Fig. 5A). In fact, the 8×CCG reporter has slightly higher expression than the 8×CCA control reporter at high initiation rate (bottom vs. top panel, *ΔASC1* in Fig. 5A). Similarly, deletion of *HEL2* increases protein expression of stall-containing reporters by 3-fold at high initiation rate compared to the wild-type strains, but has no effect at low initiation rate or in the control reporters (*ΔHEL2* in Fig. 5A). Deletion of either *DOM34* or *LTN1* has little to no effect on protein expression from either the stall-containing or control reporters (*ΔDOM34* and *ΔLTN1* in Fig. 5A). Western blotting against the N-terminus 3×FLAG epitope qualitatively matches the above measurements of protein expression using flow cytometry (Fig. 5B). The rescue of protein expression at high initiation rates in the *ASC1* and *HEL2* deletion strains is also observed with the weaker 5×CGG stalls (Fig. S4A).

We then used our pooled sequencing-based strategy to quantify the mRNA levels of our reporters (Fig. 2A) in different deletion backgrounds. Deletion of either *ASC1* or *HEL2* selectively and completely rescues the decreased mRNA levels of the 8×CCG stall-containing reporters at high initiation rates (*ΔASC1* and *ΔHEL2* in Fig. 5C). By contrast, deletion of neither *DOM34* nor *LTN1* has any effect on the decreased mRNA levels at high initiation rates (*ΔDOM34* and *ΔLTN1* in Fig. 5C).

Finally, we checked whether the observed rescue of protein expression in the *ASC1* and *HEL2* deletion strains can be reversed by constitutive expression of the respective proteins in *trans* from the *HIS3* locus (Fig. 5D). Complementing by the wild type Asc1 protein fully reverses the protein expression rescue at high initiation rates from 8×CCG stall-containing mRNAs in *ΔASC1* strains (WT, Fig. 5D, lower panel). This reversal is abrogated either partially or completely with Asc1 mutants that are defective in translational arrest ^56^ (16HNG18AAA, 38RDKAAA and 85WDKAAA, Fig. 5D, lower panel). Similarly, complementing by the wild type Hel2 protein partially reverses the protein expression rescue at high initiation rates from stall-containing mRNAs in *ΔHEL2* strains (WT, Fig. 5D, upper panel). This reversal is not observed with Hel2 mutants that cannot bind the interacting E2 enzyme Ubc4 ^53^ (C64A, C67A and ΔRING, Fig. 5D, upper panel). In contrast to the 8×CCG stall reporters, complementing with various Asc1 and Hel2 mutants had little effect on protein expression from the 8×CCA-containing control reporters (Fig. S4B).

Based on the above measurements, we conclude that Asc1 and Hel2, which have been previously associated with ribosome collision-stimulated quality control ^24, 30, 31^, are necessary to reduce protein expression and mRNA stability at high initiation rates from stall-containing mRNAs. By contrast, neither Dom34 nor Ltn1 have a role in regulating gene expression at high initiation rates.

### 6. Endogenous mRNAs with stall sequences show signatures of inefficient translation initiation

Since stall-inducing polybasic tracts decrease protein expression and mRNA stability at high initiation rates in our reporter-based experiments, do they also shape the translation of endogenous *S. cerevisiae* mRNAs? *S. cerevisiae* protein-coding sequences that contain stretches of 6 or more lysine and arginine codons within a 10-codon window lead to ribosome stalling as measured using ribosome profiling^16^. Similarly, tandem repeats of 2 or more prolines also induce ribosome pausing ^33, 34^. Over 1250 *S. cerevisiae* protein-coding sequences contain either 10-codon stretches with at least 6 lysine and arginine codons or 10-codon stretches with at least 6 proline codons. Using a high quality *S. cerevisiae* ribosome profiling dataset ^57^, we recapitulated previous observations ^16^ of increased ribosome density about 8 codons into the stall (Fig. 6A), while we did not observe a similar increase around control sequences that are enriched for glutamate or aspartate codons (Fig. S5).

**Figure 6.**
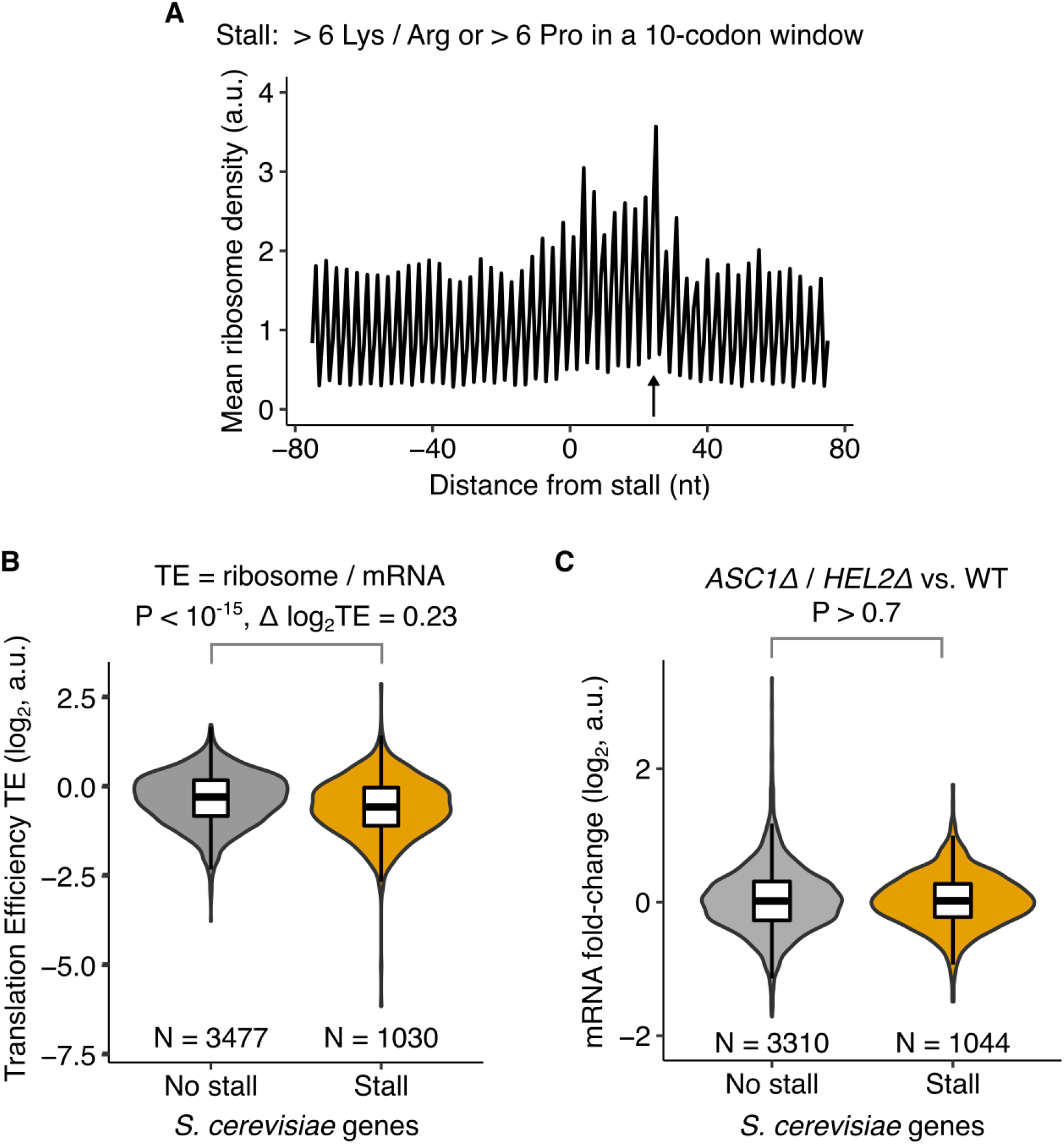
Endogenous mRNAs with stall sequences show signatures of inefficient translation initiation. **(A)** Mean ribosome density around stall-encoding sequences in *S. cerevisiae* mRNAs using data from Weinberg et al ^57^. Stall-encoding sequences are defined as 10-codon windows that have either a minimum of 6 lysine and arginine codons or a minimum of 6 proline codons. The first nucleotide of the 10-codon window is at distance 1 nt. 1251 *S. cerevisiae* mRNAs have at least one stall-encoding sequence. The ribosome density is normalized within the window around each stall-encoding sequence before calculating the mean across all sequences. Arrow indicates peak in ribosome density at +24 nt that is consistent with Brandman et al. ^16^. **(B)** Translation efficiency of *S. cerevisiae* mRNAs that either contain or do not contain stall-encoding sequences. Translation efficiency (TE) is defined as the normalized ratio of ribosome footprint counts to total mRNA counts as measured by Weinberg et al ^57^. **(C)** Fold-change in mRNA levels between both *ΔHEL2* and *ΔASC1* strains and wild-type strain. *ΔHEL2* and *ΔASC1* strains were treated as replicates in this analysis to identify mRNAs that are consistently up-regulated in both strains. mRNA levels were measured by Sitron et al. ^52^. Box plots in (B) and (C) shows mean and standard deviation within each gene group. Violin plot shows the gene density at each Y-axis value. P-values are calculated using the two-sided Wilcoxon rank-sum test.

The widespread nature of stall-encoding sequences in endogenous *S. cerevisiae* mRNAs suggests two distinct possibilities in light of the collision-stimulated decrease in gene expression uncovered in our work. First, an endogenous mRNA containing a stall could have a sufficiently high initiation rate such that it is constantly subject to collision-stimulated quality control. Indeed such a regulatory mechanism was proposed for *RQC1* which encodes one of the components of the RQC complex ^16^. Alternatively, a stall might be present due to functional requirements on the protein that are unrelated to collision-stimulated quality control. In this case, the presence of the stall might necessitate that the corresponding mRNA have sufficiently low initiation rate to prevent collision-stimulated decrease in mRNA stability and protein expression.

To distinguish between the above two possibilities, we tested whether stall-containing mRNAs have significant differences in initiation rate compared to other endogenous mRNAs in *S. cerevisiae*. We used translation efficiency (TE) as quantified by the ratio of ribosome footprints to mRNA abundance as a proxy for initiation rate ^57^. Strikingly, we find that the TE of *S. cerevisiae* mRNAs containing stall-encoding sequences (as defined above, N = 1030) is significantly lower than that of mRNAs (N = 3477) that do not contain such sequences (P < 10^−15^, Wilcoxon 2-sided test, Δ log_2_ TE = 0.23, Fig. 6B). A potential confounding factor in this analysis is that the stall-encoding sequences cause an increase in local ribosome occupancy on the corresponding mRNA. However, this increase will result in a higher apparent TE, thereby lowering the measured difference between stall-containing and control mRNAs.

Based on our observation that stall-encoding *S. cerevisiae* mRNAs have lower TE, we hypothesize that these mRNAs have been evolutionarily selected for inefficient initiation to avoid collision-stimulated reduction in protein expression and mRNA stability. This hypothesis predicts that deletion of *HEL2* or *ASC1* should have no effect on the levels of these endogenous mRNAs, unlike the increase in levels of our reporter mRNAs at high initiation rates (Fig. 5B). To verify this prediction, we compared the levels of *S. cerevisiae* mRNAs between *ΔHEL2* or *ΔASC1* strains and wild-type strains, as measured previously by RNA-seq ^52^. Since deletion of these factors can have indirect and pleiotropic effects unrelated to quality control ^58^, we looked for consistent differential regulation of mRNA levels in both *ΔHEL2* and *ΔASC1* strains compared to the wild-type strain. We find that on average, stall-containing *S. cerevisiae* mRNAs do not show a significantly higher upregulation compared to control mRNAs in the *ΔHEL2* and *ΔASC1* strains (P > 0.7, Wilcoxon 2-sided test, Fig. 6C). This observation is consistent with our hypothesis that stall-containing *S. cerevisiae* mRNAs have been evolutionary selected to avoid collision-stimulated reduction in gene expression.

## Discussion

In this work, we find that efficient translation initiation can decrease the protein expression and stability of certain eukaryotic mRNAs that undergo elongation stalls. To our knowledge, this is the first demonstration of an inverse relation between initiation rate and gene expression from a eukaryotic mRNA. While this observation is seemingly at odds with previous studies on normal eukaryotic mRNAs ^2–4^, our computational modeling shows that it arises from the interplay between normal translation and collision-stimulated quality control at elongation stalls.

Recent studies have implicated ribosome collisions in triggering endonucleolytic mRNA cleavage and ribosomal ubiquitination in eukaryotes ^24, 30, 31^. But these studies did not delineate the effect of ribosome collisions on overall mRNA stability and protein expression. Our quantitative experiments reveal that ribosome collisions can decrease mRNA stability and protein expression at high initiation rates. In addition, our computational modeling illuminates a number of kinetic constraints that shape the influence of ribosome collisions on gene expression.

Contrary to our intuitive expectation, abortive termination by leading and trailing ribosomes at collisions have qualitatively distinct effects on protein expression (CSAT vs. CAT models, Fig. 3B). Specifically, the observed decrease in protein expression at high initiation rates (Fig. 1C) can be explained only if the leading ribosome in a collision undergoes abortive termination, after accounting for changes in mRNA levels (Fig. 2C). Further, the decrease in protein expression at high initiation rates due to abortive termination requires ribosomes to stall repeatedly (Fig. 3C). This observation suggests that ribosomes stall at multiple locations within each poly-basic tract, which then leads to abortive termination. This is in contrast to elongation stalls due to amino acid starvation in bacteria, where a single stall-inducing codon is sufficient to trigger ribosome collisions, but does not cause a reduction in protein expression at high initiation rates ^42^. It will be interesting to test whether a reduction in protein expression due to polyA-mediated stalls at high initiation rates is also observed in mammalian cells ^50, 51, 53^.

Our work reveals that decrease in mRNA stability at high initiation rates is a general consequence of competition between endonucleolytic mRNA cleavage and canonical mRNA decay. Notably, we observe such a decrease in simulations of both collision-independent and collision-stimulated models of mRNA cleavage (SEC vs. CSEC models, Fig. 4B). However, the decrease in protein expression at high initiation rates occurs only when ribosome collisions stimulate endonucleolytic mRNA cleavage (SEC vs. CSEC models, Fig. 4C). Simms et al. ^24^ observed endonucleolytic mRNA cleavage only 100 nt or more 5′ to the stall site, which led them to propose that a queue of several stacked ribosomes might form an oligomer to trigger endonucleolytic mRNA cleavage. While such a model is theoretically possible, our measured decrease in protein and mRNA levels at high initiation rates can be explained by our simpler models of collision-stimulated quality control that requires sensing of only the disome interface, and is consistent with recent structural work ^30, 31^.

Our measurements of normal mRNA levels (Fig. 2B) do not support models of simple kinetic competition between translating ribosomes and canonical mRNA decay factors in *S. cerevisiae*. For example, our data is inconsistent with a model where regions of the mRNA that are not covered by ribosomes are subject to random endonucleolytic cleavage ^59^. Any competition between translation and canonical mRNA decay must occur prior to start codon recognition such as during cap binding by translation initiation factors^2^.

Our work reveals that the widely used traffic jam model ^27, 28^ (Fig. 3B) does not accurately capture ribosome dynamics on stall-containing eukaryotic mRNAs. Specifically, our measured decrease in protein expression at high initiation rate from stall-containing mRNAs is inconsistent with the traffic jam model. The traffic jam model also does not consider the interaction between translation and mRNA stability, which we observe experimentally. The collision-stimulated quality control models of ribosome kinetics formulated here (CSAT and CSEC models) provide an experimentally constrained starting point for modeling eukaryotic translation.

Recent studies have applied the traffic jam model to infer the extent of ribosome collisions from ribosome profiling measurements ^60–62^. These studies assume that collided ribosomes are under-represented in ribosome profiling data due to protocol-related biases against isolating multi-ribosome footprints. Our work instead suggests that collided ribosomes can be depleted from the pool of translated mRNAs because they stimulate abortive termination and mRNA decay. Thus the extent of ribosome interference inferred in these earlier studies using the traffic jam model might not accurately represent the *in vivo* frequency of ribosome collisions. Modeling studies that consider abortive termination assume that ribosomes randomly drop-off during elongation ^43, 44^, or that abortive termination due to ribosome collisions is negligible ^63^. Our work instead reveals that abortive termination occurs in response to ribosome collisions caused by elongation stalls, and that this process is highly regulated by cells using dedicated quality control factors.

Our analysis of endogenous *S. cerevisiae* mRNAs that contain stall-encoding sequences reveals that the initiation rates of these mRNAs might have been evolutionarily tuned to values low enough to escape ribosome collision-driven reduction in gene expression (Fig. 6). However, this aggregate analysis will not identify individual stall-containing mRNAs on which ribosome collision might have a regulatory role. Nor does our analysis consider mRNAs in which ribosome stalls are caused by sequences other than poly-basic tracts such as mRNA stem loops or other peptide features ^64^. A straightforward way by which ribosome collisions on individual mRNAs could play a functional role is through direct autoregulation, such as that previously described for the Rqc1 component of the RQC complex ^16^. A more intriguing possibility is suggested by the fact that poly-basic tracts occur in several mRNAs that encode components of either the translation apparatus or the ribosome biogenesis pathway. The stalls on these mRNAs could be part of a cellular feedback control that tunes the translational capacity of the cell by sensing ribosome collisions on mRNAs which encode the translation apparatus or enable its synthesis.

## Acknowledgments

We thank Rob Bradley, Alicia Darnell, Adam Geballe, Andrew Hsieh, Harmit Malik, Mark Roth, Gerry Smith, Brian Zid, and members of the Subramaniam lab for discussions and feedback on the manuscript. We thank Dan Gottschling, Sue Biggins, and Toshi Tsukiyama for plasmids. We thank Onn Brandman for sharing the RNA-Seq data from Sitron et al ^52^. This work was funded by grant R35 GM119835 from the National Institutes of Health (A.R.S). The computations described in this work were performed on the FHCRC research computing cluster.

## Author contributions

H.P. and A.R.S. designed and performed research, analyzed data, and wrote the paper. A.R.S. supervised the project and acquired funding.

## Materials and Methods

Programming code and instructions for reproducing the simulations and data analyses in this manuscript are publicly available at https:github.com/rasilab/ribosome_collisions_yeast. Our customized versions of the simulation software PySB, BioNetGen, and NFsim along with instructions for their use can be accessed from our laboratory’s Github page (https://github.com/rasilab).

### 1. Strain and plasmid construction

Tables S1 and S2 list the plasmid cloning vectors and *S. cerevisiae* strains used in this work. `DNA_ sequences.fasta` (https://github.com/rasilab/ribosome_collisions_yeast/blob/master/data/dna_sequences.fasta) contains the sequences of the reference plasmid vectors and inserts used in this work. Plasmids and strains will be shipped upon request. PCR oligo sequences are available upon request.

The BY4741 *S. cerevisiae* strain background (S288C, *MATa HIS3Δ1 LEU2Δ0 MET15Δ0 URA3Δ0*, Thermo Fisher) was used for all experiments in this study. The pHPSC16 vector containing *mKate2* and the *LEU2* marker was integrated as a single copy at the *HO* locus. The resulting strain (scHP15) was used as a parent for inserting all *PGK1-YFP* reporters and for deleting quality control-associated genes. The p41894 vector and its derivatives containing *PGK1-YFP* variants and the *URA3* marker were inserted as a single copy at a ChrI intergenic locus. The pSB2273 vector containing *HEL2* or *ASC1* variants and the *HIS3* marker were inserted as a single copy at the *HIS3* locus. Integration into the *S. cerevisiae* genome was performed by transforming 0.5–2 *μg* of the linearized (NotI or PmeI restriction) plasmid vector according to the LiAc/SS carrier DNA/PEG method ^65^. Single yeast colonies were selected on synthetic complete (SC) agar plates lacking either leucine, uracil, or histidine.

*LTN1, DOM34, HEL2*, or *ASC1* were deleted from the *S. cerevisiae* genome using PCR-mediated gene disruption ^66^. Deletion cassettes with flanking homology arms are provided in `DNA_sequences.fasta`. Primers for amplifying *KAN* or *NAT* resistance markers with 40–50 bp homology arms were designed using the sequences from the *S. cerevisiae* genome deletion project webpage (http://www-sequence.stanford.edu/group/yeast_deletion_project/downloads.html).

All plasmids were cloned by restricting the parent vectors and inserting 2–4 PCR-amplified fragments with 20–30 bp homology arms using isothermal assembly ^67^. The inserted sequences were confirmed by Sanger sequencing. The 5′ UTR mutations for varying initiation rate and the ribosome stall/control repeats were introduced into the PCR primers used for amplifying inserts prior to isothermal assembly. These 5’UTR and insert sequences are provided in `DNA_sequences.fasta`. Standard molecular biology procedures ^68^ were followed for all other steps of plasmid cloning.

*3×FLAG-PGK1-YFP* reporter genes were inserted into the p41894 vector ^69^ between the *GPD* (*TDH3*) promoter + 5′ UTR and the *CYC1* 3′ UTR + transcriptional terminator using the XbaI and XhoI restriction sites. The sequence around the XbaI site of p41894 was modified to retain the native 5′ UTR sequence of *TDH3* while mutating the −12 to −7 nt from ATG to XbaI (TAAACA to TCTAGA). *PGK1\** protein coding sequence was generated from wild-type *S.cerevisiae PGK1* by introducing 5×AGA repeats at the five locations used in a previous study ^11^. The *S. cerevisiae*-optimized *YFP* (mCitrineV163A variant) sequence was taken from a previous work ^70^, and was fused with *PGK1* or *PGK1\** with an intermediate BamHI site. The sequence of the resulting *p41894-3×FLAG-PGK1*-YFP* vector, pHPSC57, is provided in `DNA_sequences.fasta`. The *3×FLAG-PGK1-YFP* insert sequence between XbaI and XhoI of pHPSC417 is also provided.

The *HO3-LEU2-pGPD-mKate2-Cyc1t* vector was constructed by inserting two 400 bp sequences from the *HO* locus of the *S. cerevisiae* genome into a pUC19 backbone. The *LEU2* gene from pRS315 ^71^ and a *GPDp-mKate2-CYC1t* cassette were inserted between the two 400 bp *HO* sequences. The *mKate2* coding sequence was taken from previous work ^72^ and was synonymously mutated to the most frequent codon for each amino acid in the *S. cerevisiae* genome. The *GPDp-Cyc1t* flanking regions are identical to the ones in our reporter vector, described above. The sequence of the resulting vector, pHPSC16, is provided in `DNA_sequences.fasta`.

The *HEL2* and *ASC1* wild-type or mutant coding sequences were inserted into pSB2273 between the *GPD* promoter and the *ADH1* terminator using between the XhoI and BamHI restriction sites. The *HEL2* coding sequence was PCR-amplified from the *S. cerevisiae* genome while the *ASC1* coding sequence was PCR-amplified from *S. cerevisiae* cDNA to exclude the intron. Sequence of the parent vector, pSB2273, is provided in `DNA_sequences.fasta`. All the *HEL2* and *ASC1* insert sequences cloned into pSB2273 are also provided.

For creating the barcoded reporter strains for mRNA measurements, each of the *PGK1-YFP* reporter coding sequences was PCR-amplified separately with 4 distinct barcodes in the 3′ UTR. The ATG → CTG control reporters were amplified with 3 distinct barcodes. The PCR products were separately inserted into pHPSC57 between the XbaI and XhoI restriction sites using isothermal assembly. The isothermal assembly products were pooled and transformed at high efficiency into electro-competent *E. coli*. After overnight selection on LB-agar plates with Carbenicillin, the bacterial lawn was scraped to extract the pooled plasmids. The pooled plasmids were integrated into either scHP15 wild-type strain or one of the four deletion strain backgrounds at the ChrI intergenic locus using the same transformation protocol as above. A minimum of 500 *S. cerevisiae* colonies were pooled for each strain and stored as glycerol stocks. The pooled reporter plasmids and their respective barcodes are provided as part of the data analysis code at https://github.com/rasilab/ribosome_collisions_yeast#high-throughput-sequencing.

### 2. Flow cytometry

3–8 single *S. cerevisiae* colonies from transformation were inoculated into separate wells of 96-well plates containing 150 *μ*l of yeast extract-peptone with 2% dextrose (YEPD) medium in each well and grown overnight for 16 hours at 30°C with shaking at 800 rpm. The saturated cultures were diluted 100-fold into 150*μ*l of fresh YEPD medium. After growing for 6 hours under the same condition as overnight, the plates were placed on ice, and analyzed using the 96-well attachment of a flow cytometer (BD FACS Aria or Symphony). Forward scatter (FSC), side scatter (SSC), YFP fluorescence (FITC) and mKate2 fluorescence (PE.Texas.Red) were measured for 10,000 cells in each well. The resulting data in individual `.fcs` files for each well was combined into a single tab-delimited text file. Mean YFP and mKate2 fluorescence in each well was calculated, and the YFP background from the scHP15 strain and the mKate2 background from the BY4741 strain were subtracted. The ratio of the background-subtracted YFP to mKate2 signal was normalized by that of a standard internal control strain. The average and standard error of this ratio was calculated over all transformant replicate wells and is shown in Fig. 1 and 5. Analysis code for reproducing the figures from raw flow cytometry data is available at https:github.com/rasilab/ribosome_collisions_yeast. 2–5 outliers among several hundred wells that were not fluorescent were discarded in some experiments, and are indicated as such in the analysis code for that experiment. The mKate2 channel measurement was not steady during the flow cytometry measurement in Fig. 5D, and hence the YFP fluorescence for this experiment is plotted without normalizing by mKate2 signal. In general, normalizing by mKate2 slightly reduced the fluctuations between biological replicates but did not have any qualitative effect in any experiment. Full analysis code for flow cytometry data is provided in https://github.com/rasilab/ribosome_collisions_yeast#flow-cytometry. The P-values in Fig. 5C are calculated using the `t.test` function in R. The analysis code for calculating these P-values are provided in https://github.com/rasilab/ribosome_collisions_yeast/blob/master/analysis/flow/hel2_asc1_mutants.md.

### 3. Deep sequencing quantification of mRNA levels

*S. cerevisiae* glycerol stocks containing pooled reporter strains (> 10^7^ cells) were thawed and grown overnight in 3 ml of YEPD at 30°C in roller drums. The saturated cultures were diluted 200-fold into 50ml of fresh YEPD medium and grown for around 4:30 hours at 30°C with shaking at 200rpm. Each culture was split into two 25ml aliquots (one for extracting RNA and the other for extracting genomic DNA) into pre-chilled 50 ml conical tubes and spun down at 3000g, 4°C, 2 min. The supernatant was removed and the cell pellets were flash frozen in a dry ice / ethanol bath and stored at –80°C.

For RNA extraction, the cell pellets were re-suspended in 400*μ*l of Trizol (15596-026, Thermo Fisher) in a 1.5 ml tube and vortexed with 500*μ*l of glass beads (G8772, Sigma) at 4°C for 15 min. RNA was extracted from the resulting lysate using the Direct-zol RNA Miniprep Kit (R2070, Zymo) following manufacturer’s instructions. For genomic DNA (gDNA) extraction, the cell pellets were processed using the YeaStar Genomic DNA Kit (D2002, Zymo) following manufacturer’s instructions.

1 *μ*g of total RNA was reverse transcribed into cDNA using SuperScript IV (18090-010, Thermo Fisher) reverse transcriptase and a primer annealing to the 3′ UTR (oMF321: ACACTCTTTCCCTACACGACGCTCTTCCGATCTGCGTGACATAACTAATTACAT) following manufacturer’s instructions. A 252 nt region surrounding the 8 nt barcode was PCR-amplified from both the cDNA and gDNA in two rounds. Round 1 PCR was carried out for 10 cycles with oMF321 and oHP139 (GTGACTGGAGTTCAGACGTGTGCTCTTCCGATCTAAGGACCCAAACGAAAAG primers using Phusion polymerase (F530, Thermo Fisher) following manufacturer’s instructions. Round 2 PCR was carried out for 10 cycles for gDNA samples and 10 or 14 cycles for cDNA samples with a common reverse primer (oAS111: AATGATACGGCGACCACCGAGATCTACACTCTTTCCCTACACGACGCTC) and indexed forward primers for pooled high-throughput sequencing of different samples (CAAGCAG AAGACGGCATACGAGATxxxxxxGTGACTGGAGTTCAGACGTGTGCTC). xxxxxx refers to the 6 nt sample barcode and is provided in a table as part of our analysis script. The pooled libraries were sequenced in a HiSeq 2000 (Illumina) in a 50 nt single end rapid run mode.

The high throughput sequencing data was obtained as demultiplexed `.fastq` files. The barcode corresponding to each reporter was identified in each 50 nt read and the barcode counts for each reporter tallied within each sample. The `log2` barcode counts for the cDNA were normalized by those of the gDNA, after applying a minimum threshold of 100 counts per barcode within each sample. The average and standard error of this log2 ratio was calculated over all barcodes that crossed the 100 count threshold. The average `log2` ratio was median-normalized within each set of 5′ UTR variants for a given coding sequence and sample, and is shown in Fig. 2 and 5. Analysis code for reproducing the figures from raw high-throughput sequencing data is available at https://github.com/rasilab/ribosome_collisions_yeast#high-throughput-sequencing.

### 4. Western blotting

Overnight cultures were grown from single *S. cerevisiae* colonies in 3 ml of YEPD at 30°C in roller drums. The saturated cultures were diluted 100-fold into 3 ml of fresh YEPD medium and grown for 5 hours at 30°C in roller drums. The cultures were quickly chilled in an ice-water bath and spun down in 1.5 ml tubes. The cell pellet was washed in 500 *μ*l of water and incubated in 200 *μ*l of 0.1M NaOH for 5 min at room temperature. The pellet was re-suspended in 50 *μ*l of 1X Laemmli buffer and western blots were performed using standard molecular biology procedures ^68^. The anti-FLAG antibody (F1804, Sigma) and the anti-H3 antibody (ab1791, Abcam) were visualized using IRDye antibodies (926-68072 and 926-32211, Licor) on an Odyssey imager. The raw signal in each lane was quantified using ImageJ using the rectangle-select tool followed by the Analyze-Measure menu. For each lane, the FLAG signal was divided by the H3 signal, and normalized to a maximum of 10 within each blot. Un-cropped images of blots and their quantification are provided in https://github.com/rasilab/ribosome_collisions_yeast/tree/master/data/western. The normalization of lanes is done in https://github.com/rasilab/ribosome_collisions_yeast/blob/master/analysis/western/western_analysis.md.

### 5. Kinetic modeling of eukaryotic quality control

We specify our kinetic models using the `PySB` interface ^73^ to the `BioNetGena` modeling language ^74^. The resulting `Python` script tasep.py is parsed using the `BioNetGen` parser into the tasep.bngl file, and converted into the tasep.xml file for input to the agent-based stochastic simulator, `NFsim`^75^.

Below, we describe the molecules and the reactions in our model, along with the parameters and initiation conditions. A complete list of parameters in our model is provided in Table S3. The tasep.bngl file provides a compact but rigorous summary of the description below using `BioNetGen` language.

#### 5.1. Molecules

Our kinetic model of eukaryotic quality control has five molecule types: DNA, ribosome, mRNA, fully synthesized protein, and aborted protein. Below, we describe the components, their states and binding partners of each molecule type.

The DNA molecule does not have any components. It is used as a template for generating new mRNA molecules through transcription in order to compensate for the mRNA molecules degraded through canonical mRNA decay or endonucleolytic mRNA cleavage, and thus maintain a translatable mRNA pool in our simulation at steady-state.

The ribosome molecule has three components: an A site (*a*), a back / mRNA exit site (*b*), and a front / mRNA entry site (*f*). These sites do not have distinct states but serve as bonding sites. The A site can bond to the codon site on the mRNA. The back / mRNA exit site can bond to a collided trailing ribosome on the mRNA. The front / mRNA entry site can bond to a collided leading ribosome on the mRNA. The ribosome has an mRNA footprint of 10 codons in our simulation, which is the approximate size of a ribosome footprint in *S. cerevisiae* ^40^.

The mRNA molecule has the following components: cap, start region, codon sites (*c*_*i*_), backbone sites (*r*_*i*_), and poly-A tail sites (*p*_*i*_).

The cap can be either present or absent. The cap has to be present for translation initiation and be absent for initiation of 5′-3′ exonucleolytic mRNA decay. mRNAs are transcribed with caps in our model, since the kinetics of how mRNAs are synthesized and rendered translatable is not of interest in this work. mRNAs can be decapped either upon complete deadenylation of the poly-A tail ^41^, or after endonucleolytic mRNA cleavage ^54^.

The mRNA start region can be either clear or blocked. The start region has to be clear for translation initiation. Translation initiation renders the start region blocked, while elongation past a ribosome footprint distance from the start codon renders the start region clear. The inclusion of start region component in our model is not strictly necessary, but allows compact specification of start codon occlusion by initiating ribosomes. The alternative is explicit specification of the ribosome occupancy of every codon within the ribosome footprint distance from the start codon.

The mRNA codon sites do not have distinct states. They serve as bonding sites for the ribosome A-site. The number of mRNA codon sites *L*_*m*_ is specified by the length of the coding region in our simulation (650 corresponding to the *PGK1-YFP* reporter used in our experiments).

The mRNA backbone sites can be either intact, endonucleolytically cleaved, or exonucleolytically cleaved. These sites serve as substrates for co-translational endonucleolytic cleavage or exonucleolytic degradation. The number of mRNA backbone sites *L*_*m*_ is specified by the length of the coding region in our simulation.

The mRNA poly-A tail sites can be either intact or exonucleolytically cleaved. The poly-A sites serve as substrates for deadenylation during canonical mRNA decay. The number of mRNA poly-A sites *L*_*p*_ is specified by the length of the poly-A region in our simulation (set to 60).

The fully synthesized protein *P* and aborted protein *A* molecules do not have any components. They are used for tracking the number of full proteins and aborted proteins produced during the course of the simulation. An equivalent strategy will be to exclude these molecules and just track the ribosomes that undergo normal and premature termination events.

#### 5.2. Kinetic reactions

Below, we describe each type of kinetic reaction in our model of quality control during eukaryotic translation. We use a syntax similar to that of `BioNetGen`^74^ for illustrating kinetic reactions.

Transcription produces capped mRNAs without any ribosomes (Eq. 1). All mRNA backbone sites and poly-A tail sites are intact. The transcription rate constant *k*_*transcription*_ is set to 0.001*s*^−1^. The value of this parameter is arbitrary, and is chosen to maintain a pool of 1–2 translatable mRNAs at any given time in the simulation, and to produce around 1000 mRNAs during a typical simulation run for 10^6^*s*.

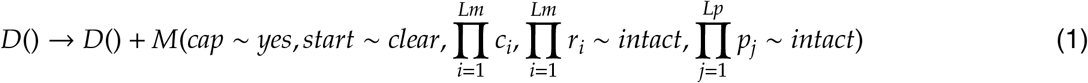

Translation initiation occurs on capped mRNAs when the start region is free, the start codon is not bonded, and the mRNA backbone at the start codon is intact (Eq. 2). It results in the start region being blocked, and the ribosome A-site being bonded to the start codon (‘!1’ in Eq. 2). The translation initiation rate *k*_*init*_ is systematically varied in our simulations. We use a default value of 0.1*s*^−1^ to match previous estimates ^38, 76^. The absolute value of this parameter is not critical in our modeling. Rather, the dimensionless ratio of the translation initiation rate to elongation rate is the important parameter in our model since it determines the frequency of ribosome collisions at the stall.

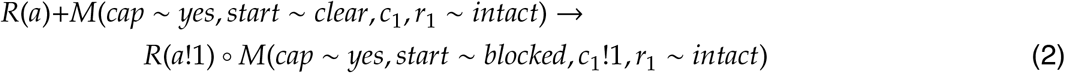

Elongation of a ribosome results in the ribosome A-site moving from codon *c*_*i*_ to codon *c*_*i*+1_ (Eq. 3). Elongation requires that there is no leading ribosome within a footprint distance at *c*_*i*+10_. Additionally, when a ribosome elongates at the 9th codon from the initiation codon, the mRNA start region is cleared for a new round of initiation (Eq. 4). Finally, elongation of ribosomes that have been hit from the back results in loss of the molecular interaction with the trailing ribosome (‘!3’ in Eq. 5). The elongation rate in all these cases, *k*_*elong*_, is set to 10*s*^−1^ to match experimental estimates ^77^.

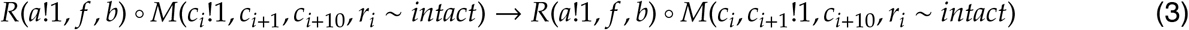

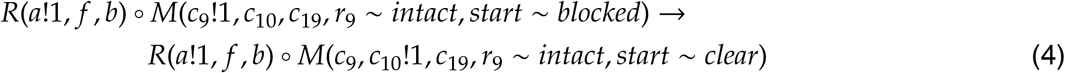

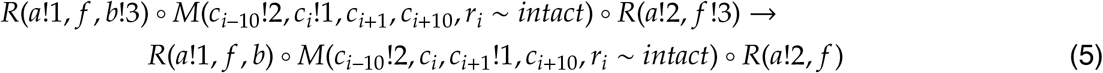

Termination results in dissociation of the ribosome-mRNA complex along with production of a protein molecule (Eq. 6). Termination of ribosomes that have been hit from the back results in loss of the molecular interaction with the trailing ribosome (Eq. 7). The termination rate, *k*_*term*_, is set to 1*s*^−1^ to be lower than the normal elongation rate of 10*s*^−1^. This choice reflects the observation that ribosome density at stop codons tends to be several fold higher than at sense codons ^34^. The absolute value of *k*_*term*_ is not important in our modeling as long as it is greater than the total rate of initiation and elongation across the mRNA.

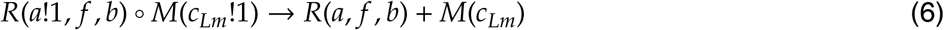

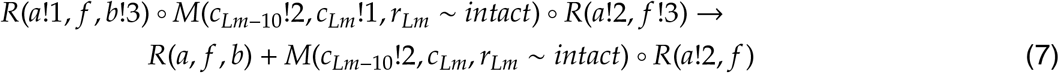

Collision between two ribosomes requires them to be separated by exactly a footprint distance on an intact mRNA, and results in a bond (‘!3’ in Eq. 8) between the *f* site of the trailing ribosome and the *b* site of the leading ribosome. The collision rate constant is set equal to the elongation rate at the A-site codon of the trailing ribosome. This choice reflects our assumption that collision occurs when the trailing ribosome tries to translocate to the next codon on the mRNA, but is sterically blocked by the leading ribosome.

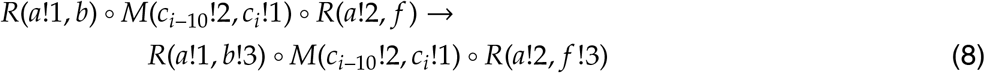

Deadenylation of the poly-A tail occurs in the 3′-5′ direction. Deadenylation at position *j* requires that the A at position *j* + 1 has been removed (Eq. 9). This constraint does not apply to the 3′ end of the poly-A tail (at position *l*_*p*_) whose removal begins the process of deadenylation (Eq. 10). We set the deadenylation rate per nucleotide *k*_*deadenylation*_ to be 0.03*s*^−1^ and the poly-A tail to be 60 nt to match previous estimates for the *PGK1* mRNA ^41^. The deadenylation rate sets the overall rate of canonical mRNA decay in our modeling. Our model of deadenylation can be refined further based on recent biochemical and genome-wide studies of poly-A tail metabolism ^14, 78^. We do not pursue this here since we do not monitor poly-A tail length and our focus is on modeling co-translational quality control.

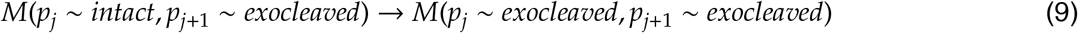

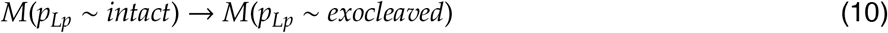

Decapping during canonical mRNA decay occurs after the poly-A tail has been fully deadenylated (Eq. 11). We set the decapping rate constant *k*_*decapping*_ to be 0.01*s*^−1^, similar in magnitude to previous work ^41^. Since we do not know the translation efficiency of deadenylated but capped mRNAs, we assume that mRNAs are initiated at their normal efficiency right until the cap is removed (Eq. 2).

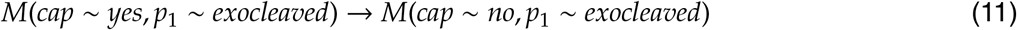

5′–3′ exonucleolysis at position *i* requires that the mRNA backbone at position *i* – 1 has been removed (Eq. 12), and that the position is not blocked by a ribosome. 5′–3′ exonucleolysis can initiate during canonical mRNA decay only after the mRNA has been decapped (Eq. 13). We set the 5′–3′ exonucleolysis rate constant *k*_*exo*_53_ to be 1*s*^−1^ ^41^, such that it is faster than the total rate of deadenylation and decapping.

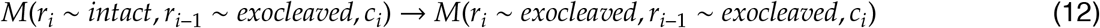

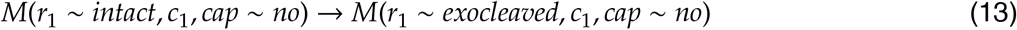

3′–5′ exonucleolysis at position *i* requires that the mRNA backbone at position *i* + 1 has been removed (Eq. 14), and that the position is not blocked by a ribosome. 3′–5′ exonucleolysis can initiate during canonical mRNA decay only after the mRNA has been fully deadenylated (Eq. 15). We set the 3′–5′ exonucleolysis rate constant *k*_*exo_*53_ to be 0. This choice is consistent with the much slower observed rate of 3′–5′ exonucleolysis in comparison with 5′–3′ exonucleolysis in *S. cerevisiae* ^41^. Further, any mRNA undergoing 3′–5′ exonucleolysis will not contribute to the pool of full length proteins or mRNA molecules, which are the experimental observables of interest in this work.

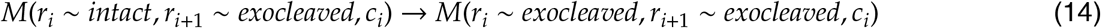

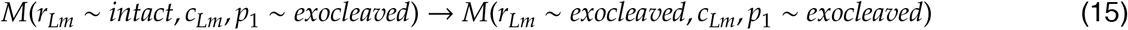

Abortive (premature) termination results in dissociation of the aborting ribosome from the mRNA. Abortive termination of a ribosome also results in the loss of molecular interaction with any collided ribosome in the front or the back. In our modeling of collision-stimulated quality control from intact mRNAs, ribosomes have different rates of abortive termination depending on whether they have undergone collision from the front (*k*_*abort_front_hit*_, Eq. 18), back (*k*_*abort_back_hit*_, Eq. 17), or both (*k*_*abort_both_hit*_, Eq. 19) sides. Ribosomes that have not undergone collisions also abortively terminate (*k*_*abort_no_hit*_, Eq. 16). The different models of abortive termination in our work (Fig. 3A) correspond to different combinations of the four types of abortive termination. In the SAT model (Eq. 8 in Fig. 3A), *k*_*abort_no_hit*_ = *k*_*abort_back_hit*_ > 0 and *k*_*abort_no_hit*_ = *k*_*abort_front_hit*_ = 0. This choice implies that collisions do not have any stimulatory effect on abortive termination, and that ribosomes that are stacked behind the leading stalled ribosome do not undergo abortive termination. In the CSAT model (Eq. 9 in Fig.3A), *k*_*abort_back_hit*_ = *k*_*abort_both_hit*_ > 0 and *k*_*abort_no_hit*_ = *k*_*abort_front_hit*_ = 0. This choice implies that collisions from back are necessary for abortive termination, but ribosomes that are stalled either on their own or due to a leading stacked ribosome do not undergo abortive termination. In the CAT model (Eq. 10 in Fig. 3A), *k*_*abort_front_hit*_ = *k*_*abort_both_hit*_ > 0 and *k*_*abort_no_hit*_ = *k*_*abort_back_hit*_ = 0. This choice implies that collisions from front are necessary for abortive termination, but ribosomes that are stalled without a leading stacked ribosome do not undergo abortive termination. The *in vivo* abortive termination rates are not known. In each of the above three models, we chose the respective non-zero abortive termination rates to be such that they would decrease protein expression appreciably due to stalling.

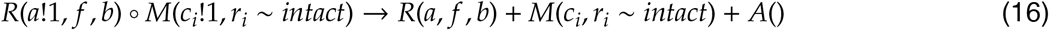

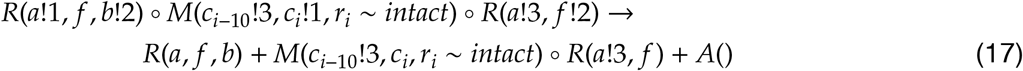

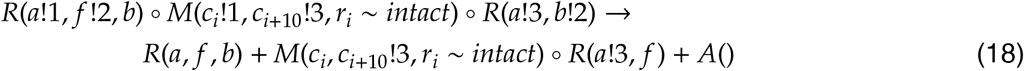

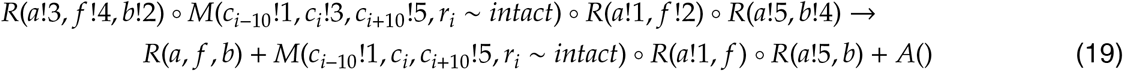

We also modeled abortive termination of ribosomes with a truncated mRNA in its A-site using the same rules as above but with *r*_*i*_ *∼endocleaved* instead of *r*_*i*_ *∼intact* This is necessary for releasing all the ribosomes that are 5′ to the endonucleolytic cleavage site. We assume that ribosomes abort with a uniformly high rate of 1*s*^−1^ from mRNAs with cleaved A-sites independent of whether they have undergone collision. This is consistent with previous *in vitro* experiments showing that Dom34/HbsI can efficiently recycle ribosomes that are stalled on a truncated mRNA ^19, 20^.

Endonucleolytic mRNA cleavage on translating ribosomes results in the mRNA being cleaved at a distance *Lc* 5′ to the A-site. We further assume that the endonucleolytic cleavage triggers immediate decapping. This assumption is consistent with increased rates of degradation of mRNAs that have undergone endonucleolytic cleavage ^54^. This assumption does not affect any of the conclusions in this work since further rounds of initiation of cleaved mRNAs will not in production of full length proteins, the observable of interest to us. In our modeling of collision-stimulated endonucleolytic cleavage, we consider different rates of endonucleolytic mRNA cleavage depending on whether ribosomes have undergone collision from the front (*k*_*cleave_front_hit*_,21), back (*k*_*cleave_back_hit*_, Eq. 22), or both (*k*_*cleave_both_hit*_, Eq. 23) sides. Ribosomes that have not undergone collisions also trigger endonucleolytic mRNA cleavage (_*kcleave_no_hit*_, Eq. 20) The different models of cotranslational endonucleolytic cleavage in our work (Fig. 4A) correspond to different combinations of the four types of endonucleolytic cleavage. In the SEC model (Eq. 11 in Fig. 4A), *k*_*cleave_no_hit*_ = *k*_*cleave_back_hit*_ > 0 and *k*_*cleave_f* *ront_hit*_ = *k*_*cleave_both_hit*_ = 0. This choice implies that ribosome collisions do not have any stimulatory effect on endonucleolytic mRNA cleavage, and that ribosomes that are stacked behind the leading stalled ribosome do not cause endonucleolytic mRNA cleavage. In the CSEC model (Eq. 12 in Fig. 4A), *k*_*cleave_back_hit*_ = *k*_*cleave_both_hit*_ > 0 and *k*_*cleave_no_hit*_ = *k*_*cleave_front_hit*_ = 0. This choice implies that collisions from back are necessary for stimulating endonucleolytic mRNA cleavage, but ribosomes that are stalled either on their own or due to a leading stacked ribosome do not result in endonucleolytic cleavage. Unlike the case of abortive termination, we do not separately model the effect of the trailing vs leading ribosome in a collision stimulating endonucleolytic cleavage, since both these models will result in a cleaved mRNA.

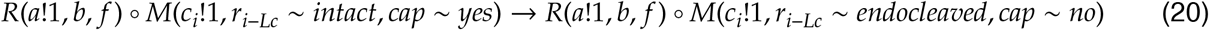

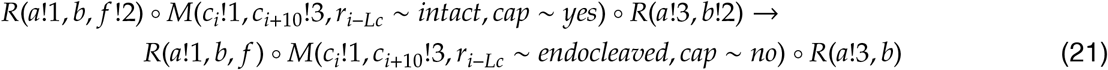

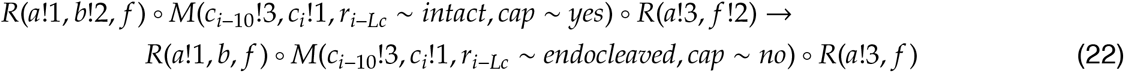

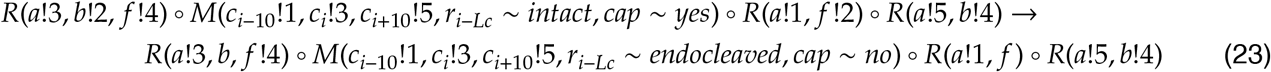

We set *Lc* = 10in our simulations even though experiments indicate that endonucleolytic mRNA cleavage occurs 14-15nt behind the stalled ribosome ^47, 48^. This choice of *Lc* to be equal to the ribosome footprint size simplifies the specification of reaction rules, but does not affect the predictions for either protein output or mRNA lifetime. This can be set to the more physiological value of 5 in case the model predictions for cleaved fragments are examined (such as when comparing against short ribosome footprints ^47^). The *in vivo* endonucleolytic cleavage rates are not known. In the above two models, we choose the respective non-zero endonucleolytic mRNA cleavage rates to be such that they would decrease mRNA lifetimes appreciably due to stalling.

#### 5.3. Simulation of quality control model

All scripts below are available at https://github.com/rasilab/ribosome_collisions_yeast/modeling. The `Python` script `choose_simulation_parameters.py` specifies the combination of parameters that we simulate using the `tasep.py` model. The `Python` script `run_simulation.py` imports the `PySB` model specified in `tasep.py`, and then invokes the `NFsim` simulation with a specific parameter combination determined by a command-line argument. The script `run_simulation.py` also invokes the `R` script `get_mrna_lifetime _and_psr.R` that calculates the mean and standard deviation of mRNA lifetimes and protein synthesis rate in each simulation. Protein synthesis rate is defined as the ratio of the number of full proteins produced during the simulation to the biological time for which the simulation was run (typically 10^6^*s*). The lifetime of each mRNA molecule in the simulation is defined as the interval from the time of transcription to the initiation of 5′–3′ exonucleolysis of the mRNA. Since the transcription rate constant was set to 1 mRNA molecule every 10^3^*s* in our simulations, ∼ 1000 mRNA molecules are followed from birth to death in each simulation lasting for 10^6^ seconds. In the case of simulations in which mRNAs were not degraded (Fig. 3C), the transcription rate is set to zero and a single initial mRNA is tracked during the course of the full simulation. All the simulation steps are implemented as `Snakemake` workfows^79^ in the `Snakefile` script.

### 6. Analysis of FACS-Seq data from Dvir et al. PNAS 2013

Supplementary Table S1 is downloaded from http://www.pnas.org/lookup/suppl/doi:10.1073/pnas.1222534110/-/DCSupplemental/sd01.xlsx. The `exp` column in this table is averaged over all `sequence_ variant` rows that share the same last 4 nucleotides. The mean and standard error of this average is shown in Fig. S1C.

### 7. Ribosome proiling analysis

Raw `.fastq` files from a previous study ^57^ were downloaded for the SRA experiments, `SRX391789` (RNA-seq) and `SRX391790` (Ribo-seq). The adapters were removed using `cutadapt` ^80^ with the arguments `--cut=8 --adapter=TCGTATGCCGTCTTCTGCTTG --minimum-length=15`. The trimmed reads were first depleted of rRNA contaminant reads and then aligned to the *S. cerevisiae* transcriptome and genome (`sacCer3`) using `tophat 2.0.14` ^81^ with the arguments `--bowtie1 --num-threads 8 --no-novel-juncs --library-type fr-unstranded --keep-tmp --read-mismatches 2`. The resulting `.bam` files were sorted and indexed using `samtools` ^82^. The aligned reads were assigned to the 13 nt from the 5′ end and this was used to calculate coverage across the `sacCer3` genome.

To identify stall sequences, the protein-coding sequences not marked as `Dubious` in the `sacCer3` genome (annotation set: `saccharomyces_cerevisiae_R64-1-1`) were scanned for 10 codon windows that contained at least 6 lysine and arginine codons, or 6 proline codons. The control sequences were similarly identified by scanning for windows that contained at least 6 glutamate and aspartate codons.

The Ribo-seq coverage in the 150 nt window each stall or control sequence was normalized by the mean coverage within that window. This normalized coverage was then averaged at each position in the window across all stall or control sequences, and shown in Fig. 6A, S5.

To calculate translation efficiency (TE) of *S. cerevisiae* genes, the Ribo-seq and RNA-seq `rpkm` values for each gene were downloaded from ftp://ftp.ncbi.nlm.nih.gov/geo/series/GSE75nnn/GSE75897/suppl/. TE for each gene was calculated as the ratio of the Ribo-seq `rpkm` to RNA-seq `rpkm`. Only genes with a minimum of 5 `rpkm` in each sample were used for the TE calculation.

### 8. RNA-seq analysis

Raw. `fastq` files from a previous study ^52^ were used for differential gene expression analysis between *ΔHEL2, ΔASC1*, and wild-type strains. Only RNA-seq data from strains containing control reporters were used for the analysis. Any ambiguous terminal nucleotides were trimmed using cutadapt ^80^ with the argument `--trim-n`. The trimmed reads were first depleted of rRNA contaminant reads and then aligned to the *S. cerevisiae* genome `(sacCer3)` using `bowtie 1.1.1` ^83^ with the arguments `-v 2 --un /dev/null --threads 8 --sam`. The resulting. `sam` files were converted to the binary .bam format, sorted, and indexed using `samtools` ^82^. The alignments were used to calculate the number of reads aligning to each transcript in the `TxDb.Scerevisiae.UCSC.sacCer3.sgdGene` R package using the `findOverlaps` function from `GenomicAlignments` R package ^84^. Transcripts that had a minimum of 100 alignments in each sample were used for differential gene expression analysis using `DESeq2` with default parameters ^85^. The *ΔHEL2* and *ΔASC1* samples were treated as replicates for `DESeq2` input and compared to the wild-type strain. The `log2FoldChange` output from the `results` function in `DESeq2` was used for plotting Fig. 6C.

## Supplementary Tables

**Figure S1.**
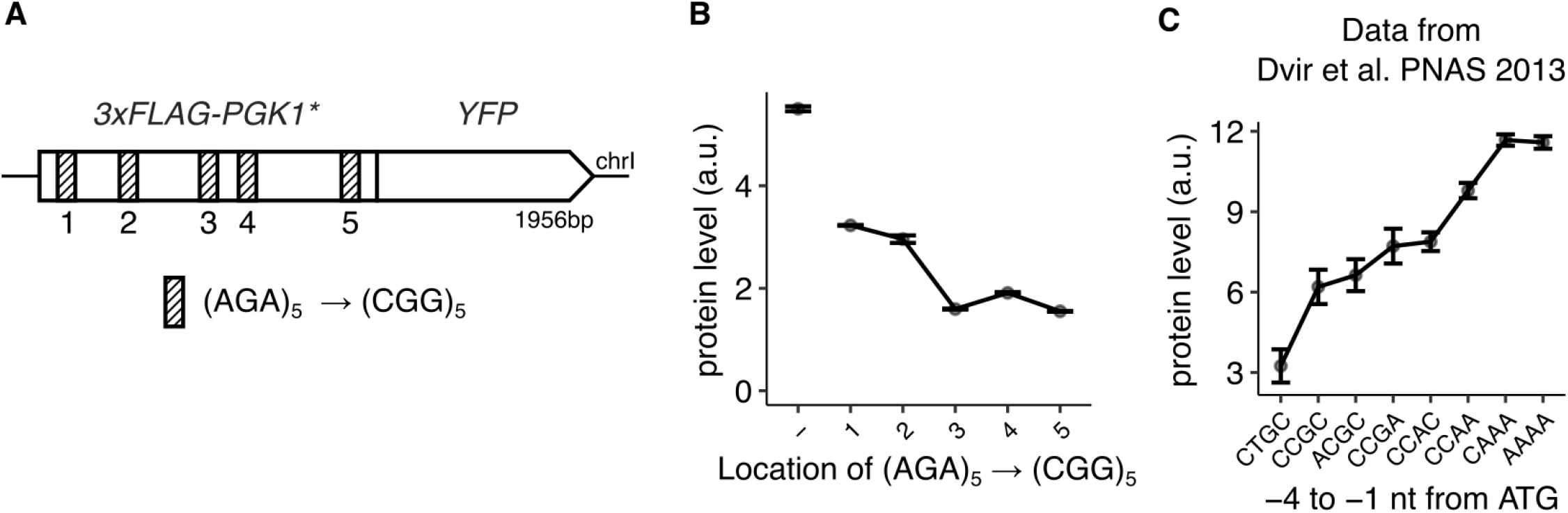
Effect of stall location on protein expression. **(A)** Schematic of *3×FLAG-PGK1*-YFP* reporters used in B. The hatched regions indicate the five locations where an (AGA)_5_ is inserted into *PGK1*. One of these locations is synonymously mutated to (CGG)_5_ in the constructs shown in B (along with the no mutation control). **(B)** Protein levels of *3×FLAG-PGK1*-YFP* reporters with with (CGG)_5_ inserted at one of the five locations indicated in A. The no-mutations control is shown as-.Protein levels are quantified as the mean fluorescence of 10,000 cells for each strain using flow cytometry. Error bars show standard error of the mean over 4 independent yeast transformants. **(C)** Protein levels of *YFP* library with randomized 10 nt region preceding the ATG start codon. Data from Dvir et al.^32^. Measured protein levels of all constructs with the same −4 to −1 nt preceding ATG are averaged and the error bars represent standard error of this average.

**Figure S2.**
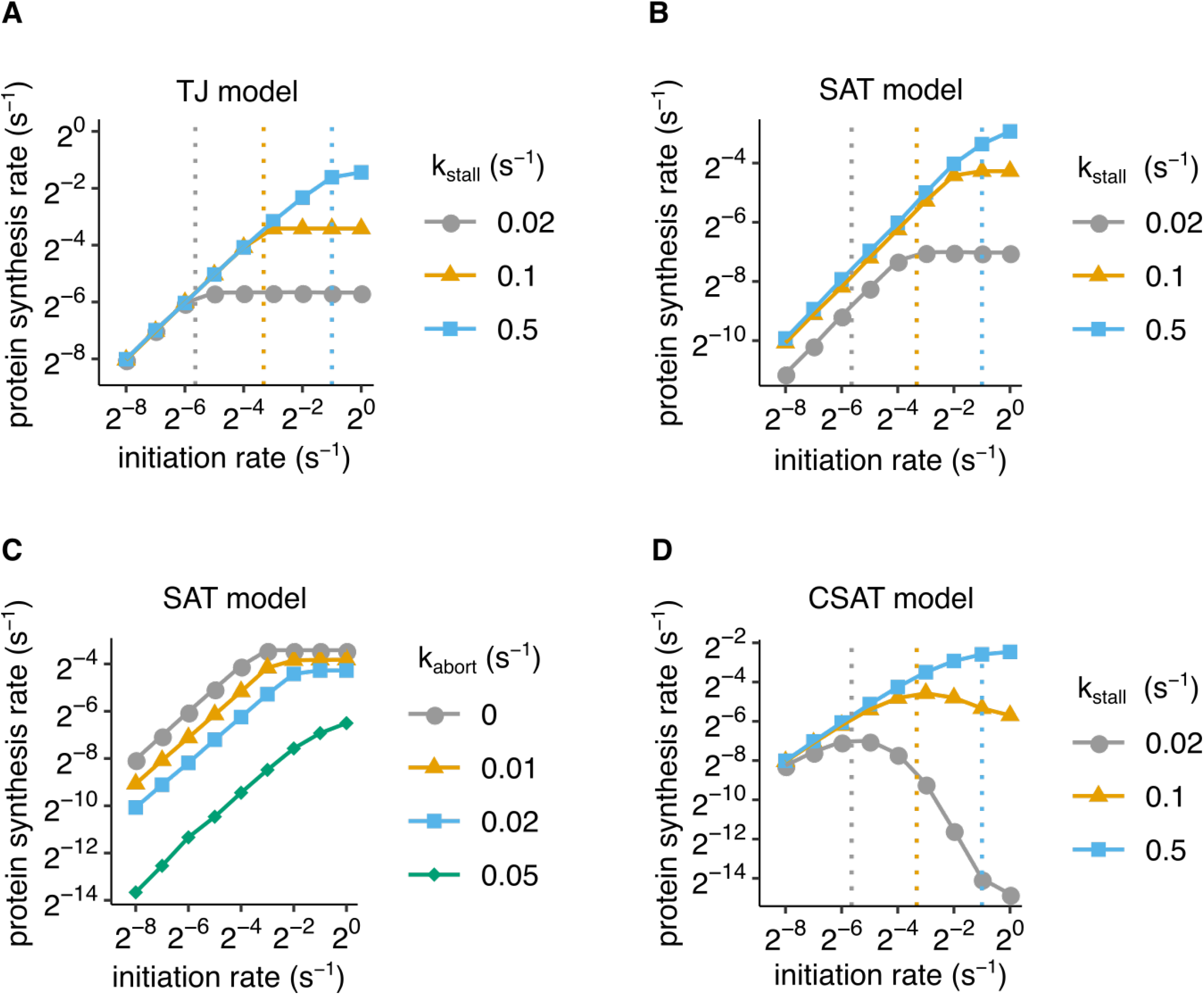
Simulated effect of elongation and abortive termination rates in the TJ, SAT, and CSAT models. (A, B, D) Protein synthesis rate as a function of initiation rate for different rates of elongation at the ribosome stall in the TJ (A), SAT (B), and CSAT (D) models. The different elongation rates at the stall are indicated graphically as vertical dashed lines for comparison with initiation rate. **(C)** Protein synthesis rate as a function of initiation rate for different rates of abortive termination at the ribosome stall in the SAT model. The mRNA is 650 codons long, and the stall is encoded by six slow translated codons located after 400 codons from the start. All other model parameters are listed in Table S3.

**Figure S3.**
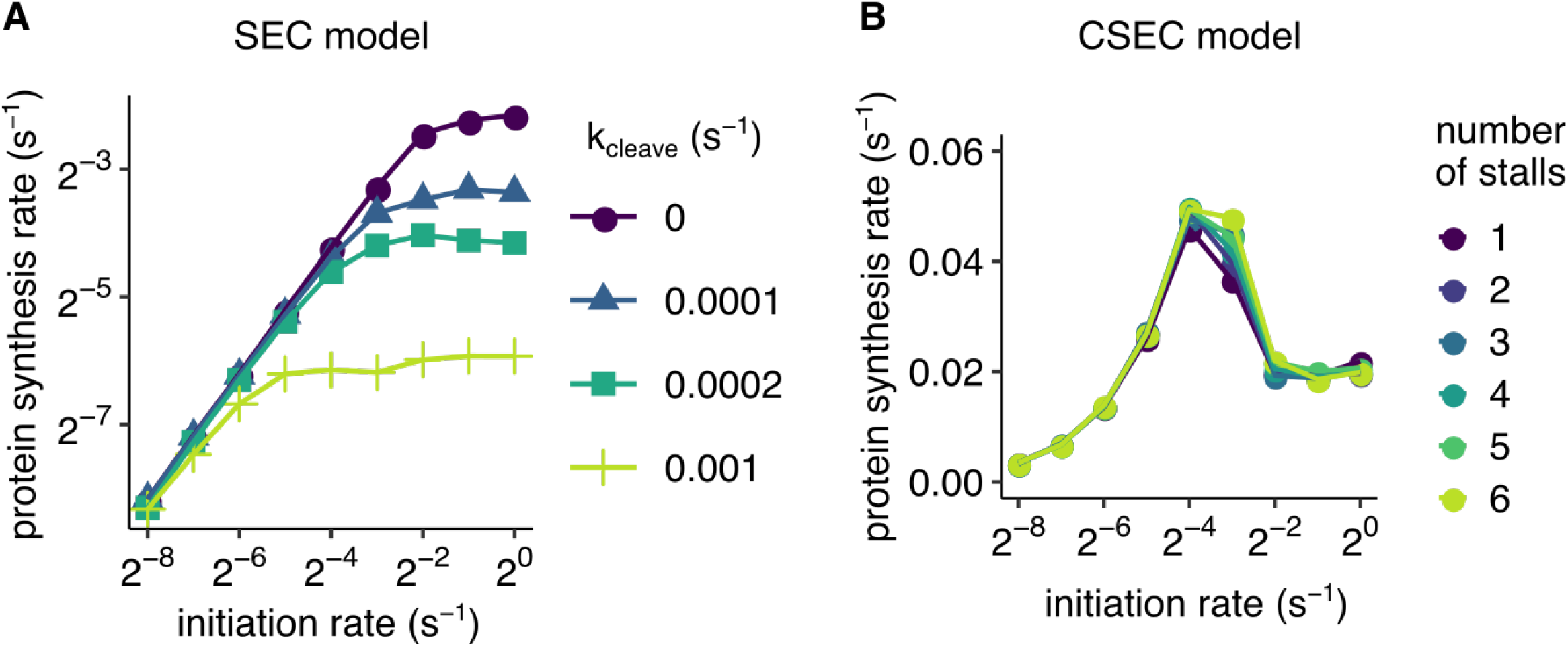
Simulated effect of cleavage rate and number of stalls in the SEC and CSEC models. **(A)** Protein synthesis rate as a function of initiation rate for different rates of co-translational endonucleolytic cleavage in the SEC model. **(B)** Protein synthesis rate as a function of initiation rate for different number of codons encoding the ribosome stall in the CSEC model. The mRNA is 650 codons long, and the stall is encoded by six slow translated codons located after 400 codons from the start in (A). All other model parameters are listed in Table S3.

**Figure S4.**
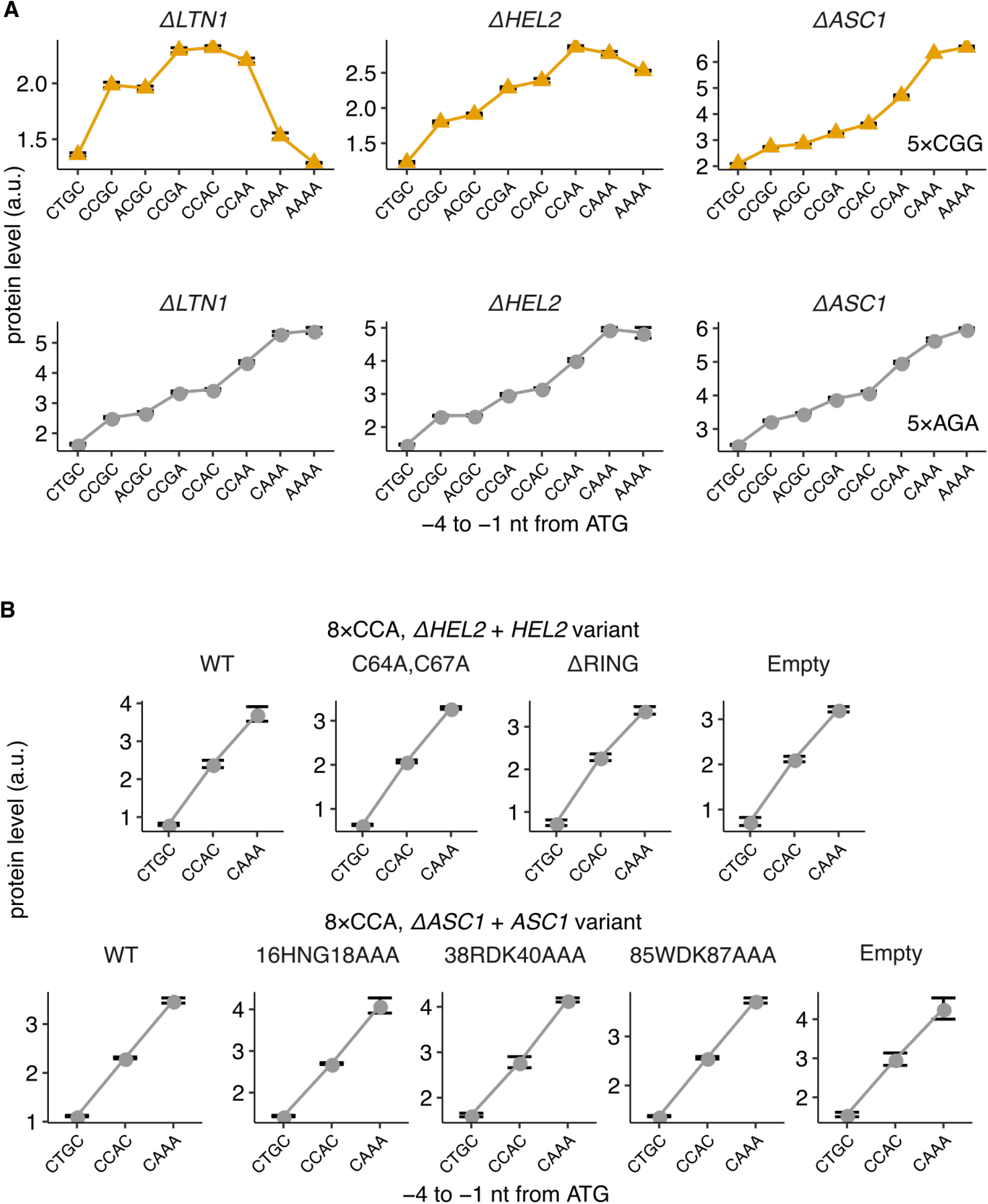
Repressive effect of high initiation rate on gene expression requires Hel2/ZNF598 and Asc1/RACK1. **(A)** Protein levels of *3–FLAG-PGK1*-YFP* reporters (see Fig. 1A) with varying initiation rates and with stall (5–CGG) or control (5–AGA) repeats. The reporters were integrated into the genome of isogenic strains with individual full deletions of *LTN1, HEL2*, or *ASC1*. **(B)** Protein levels of the 8–CCA control reporter expressed in either *ΔHEL2* (top) or *ΔASC1* (bottom) strain and complemented with the indicated HEL2 or ASC1 variant, respectively. Error bars in (A) and (B) show standard error over 3 or 4 independent transformants. The *ΔASC1*-CAAA-5–CGG variant alone has a single transformant.

**Figure S5.**
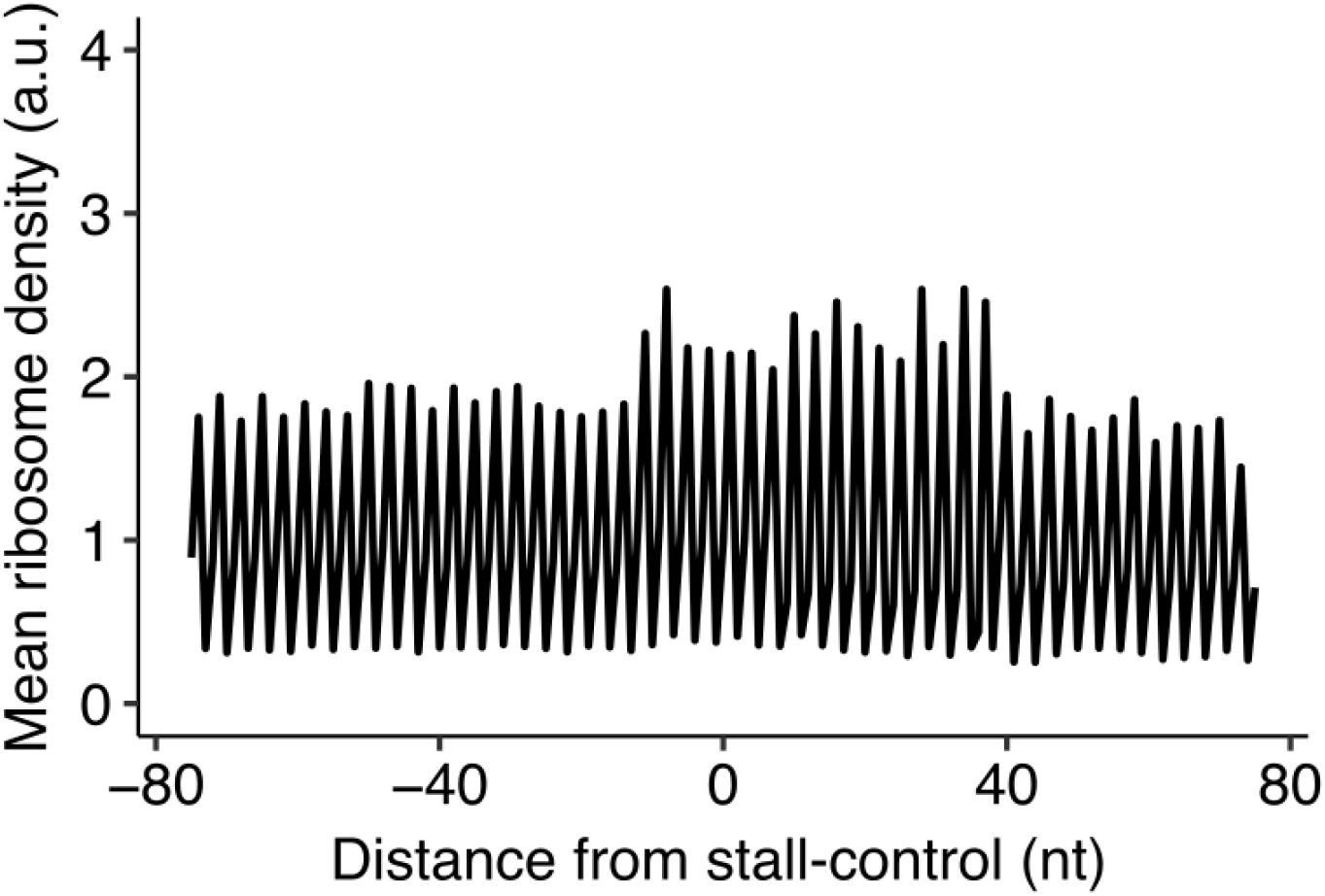
Ribosome density around control sequences does not show an increase. Mean ribosome density around control sequences using data from Weinberg et al ^57^. Control sequences are defined as 10-codon windows that have a minimum of 6 glutamate or aspartate codons. A total of 1552 *S. cerevisiae* mRNAs have at least one such control sequence. The ribosome density is normalized within the window around each control sequence before calculating the mean across all sequences.

**Table S1:** List of plasmids used for this study

| Plasmid | Genotype | Figure | Source + Comment |
| --- | --- | --- | --- |
| p41894 | chrI-URA3-pGPD-ccdB-Cyc1t | parent | Addgene 41894, single copy integration vector for ChrI safe harbor locus with URA3 selection marker |
| pHPSC16 | HO3-LEU2-pGPD-mKate2-Cyc1t | parent | This work, single copy integration vector for HO locus with LEU2 selection marker |
| pSB2273 | HIS3-pGPD-osT-Adh1t | parent | Gift from Sue Biggins lab, single copy integration vector for HIS3 locus with HIS3 selection marker |
| pHPSC417 | p41894-CAAA-ATG-3xFLAG-PGK1-YFP | 1 | This work, exp133 |
| pHPSC418 | p41894-CCGC-ATG-3xFLAG-PGK1-YFP | 1 | This work, exp133 |
| pHPSC419 | p41894-CCAA-ATG-3xFLAG-PGK1-YFP | 1 | This work, exp133 |
| pHPSC420 | p41894-CCAC-ATG-3xFLAG-PGK1-YFP | 1 | This work, exp133 |
| pHPSC421 | p41894-CCGA-ATG-3xFLAG-PGK1-YFP | 1 | This work, exp133 |
| pHPSC422 | p41894-CTGC-ATG-3xFLAG-PGK1-YFP | 1 | This work, exp133 |
| pHPSC423 | p41894-AAAA-ATG-3xFLAG-PGK1-YFP | 1 | This work, exp133 |
| pHPSC424 | p41894-ACGC-ATG-3xFLAG-PGK1-YFP | 1 | This work, exp133 |
| pHPSC425 | p41894-CAAA-CTG-3xFLAG-PGK1-YFP | 1 | This work, exp133 |
| pHPSC354 | p41894-CAAA-ATG-3xFLAG-PGK1*-10xAAG-YFP | 1 | This work, exp98 |
| pHPSC355 | p41894-CCGC-ATG-3xFLAG-PGK1*-10xAAG-YFP | 1 | This work, exp98 |
| pHPSC356 | p41894-CCAA-ATG-3xFLAG-PGK1*-10xAAG-YFP | 1 | This work, exp98 |
| pHPSC357 | p41894-CCAC-ATG-3xFLAG-PGK1*-10xAAG-YFP | 1 | This work, exp98 |
| pHPSC358 | p41894-CCGA-ATG-3xFLAG-PGK1*-10xAAG-YFP | 1 | This work, exp98 |
| pHPSC359 | p41894-CTGC-ATG-3xFLAG-PGK1*-10xAAG-YFP | 1 | This work, exp98 |
| pHPSC360 | p41894-AAAA-ATG-3xFLAG-PGK1*-10xAAG-YFP | 1 | This work, exp98 |
| pHPSC361 | p41894-ACGC-ATG-3xFLAG-PGK1*-10xAAG-YFP | 1 | This work, exp98 |
| pHPSC363 | p41894-CAAA-ATG-3xFLAG-PGK1*-10xAGA-YFP | 1 | This work, exp98 |
| pHPSC364 | p41894-CCGC-ATG-3xFLAG-PGK1*-10xAGA-YFP | 1 | This work, exp98 |
| pHPSC365 | p41894-CCAA-ATG-3xFLAG-PGK1*-10xAGA-YFP | 1 | This work, exp98 |
| pHPSC366 | p41894-CCAC-ATG-3xFLAG-PGK1*-10xAGA-YFP | 1 | This work, exp98 |
| pHPSC367 | p41894-CCGA-ATG-3xFLAG-PGK1*-10xAGA-YFP | 1 | This work, exp98 |
| pHPSC368 | p41894-CTGC-ATG-3xFLAG-PGK1*-10xAGA-YFP | 1 | This work, exp98 |
| pHPSC369 | p41894-AAAA-ATG-3xFLAG-PGK1*-10xAGA-YFP | 1 | This work, exp98 |
| pHPSC370 | p41894-ACGC-ATG-3xFLAG-PGK1*-10xAGA-YFP | 1 | This work, exp98 |

| Continued from previous page |  |  |  |
| --- | --- | --- | --- |
| Plasmid | Genotype | Figure | Source + Comment |
| pHPSC314 | p41894-CAAA-ATG-3xFLAG-PGK1*-8xCCA-YFP | 1 | This work, exp93 |
| pHPSC315 | p41894-CCGC-ATG-3xFLAG-PGK1*-8xCCA-YFP | 1 | This work, exp93 |
| pHPSC316 | p41894-CCAA-ATG-3xFLAG-PGK1*-8xCCA-YFP | 1 | This work, exp93 |
| pHPSC317 | p41894-CCAC-ATG-3xFLAG-PGK1*-8xCCA-YFP | 1 | This work, exp93 |
| pHPSC318 | p41894-CCGA-ATG-3xFLAG-PGK1*-8xCCA-YFP | 1 | This work, exp93 |
| pHPSC319 | p41894-CTGC-ATG-3xFLAG-PGK1*-8xCCA-YFP | 1 | This work, exp93 |
| pHPSC320 | p41894-AAAA-ATG-3xFLAG-PGK1*-8xCCA-YFP | 1 | This work, exp93 |
| pHPSC321 | p41894-ACGC-ATG-3xFLAG-PGK1*-8xCCA-YFP | 1 | This work, exp93 |
| pHPSC323 | p41894-CAAA-ATG-3xFLAG-PGK1*-8xCCG-YFP | 1 | This work, exp93 |
| pHPSC324 | p41894-CCGC-ATG-3xFLAG-PGK1*-8xCCG-YFP | 1 | This work, exp93 |
| pHPSC325 | p41894-CCAA-ATG-3xFLAG-PGK1*-8xCCG-YFP | 1 | This work, exp93 |
| pHPSC326 | p41894-CCAC-ATG-3xFLAG-PGK1*-8xCCG-YFP | 1 | This work, exp93 |
| pHPSC327 | p41894-CCGA-ATG-3xFLAG-PGK1*-8xCCG-YFP | 1 | This work, exp93 |
| pHPSC328 | p41894-CTGC-ATG-3xFLAG-PGK1*-8xCCG-YFP | 1 | This work, exp93 |
| pHPSC329 | p41894-AAAA-ATG-3xFLAG-PGK1*-8xCCG-YFP | 1 | This work, exp93 |
| pHPSC330 | p41894-ACGC-ATG-3xFLAG-PGK1*-8xCCG-YFP | 1 | This work, exp93 |
| pHPSC57 | p41894-CAAA-ATG-3xFLAG-PGK1*-5xAGA-YFP | 1 | This work, exp60 |
| pHPSC126 | p41894-CCGC-ATG-3xFLAG-PGK1*-5xAGA-YFP | 1 | This work, exp60 |
| pHPSC158 | p41894-CCAA-ATG-3xFLAG-PGK1*-5xAGA-YFP | 1 | This work, exp60 |
| pHPSC159 | p41894-CCAC-ATG-3xFLAG-PGK1*-5xAGA-YFP | 1 | This work, exp60 |
| pHPSC160 | p41894-CCGA-ATG-3xFLAG-PGK1*-5xAGA-YFP | 1 | This work, exp60 |
| pHPSC161 | p41894-CTGC-ATG-3xFLAG-PGK1*-5xAGA-YFP | 1 | This work, exp60 |
| pHPSC162 | p41894-AAAA-ATG-3xFLAG-PGK1*-5xAGA-YFP | 1 | This work, exp60 |
| pHPSC163 | p41894-ACGC-ATG-3xFLAG-PGK1*-5xAGA-YFP | 1 | This work, exp60 |
| pHPSC72 | p41894-CAAA-ATG-3xFLAG-PGK1*-5xCGG-YFP | 1 | This work, exp60 |
| pHPSC131 | p41894-CCGC-ATG-3xFLAG-PGK1*-5xCGG-YFP | 1 | This work, exp60 |
| pHPSC188 | p41894-CCAA-ATG-3xFLAG-PGK1*-5xCGG-YFP | 1 | This work, exp60 |
| pHPSC193 | p41894-CCAC-ATG-3xFLAG-PGK1*-5xCGG-YFP | 1 | This work, exp60 |
| pHPSC168 | p41894-CCGA-ATG-3xFLAG-PGK1*-5xCGG-YFP | 1 | This work, exp60 |
| pHPSC198 | p41894-CTGC-ATG-3xFLAG-PGK1*-5xCGG-YFP | 1 | This work, exp60 |
| pHPSC203 | p41894-AAAA-ATG-3xFLAG-PGK1*-5xCGG-YFP | 1 | This work, exp60 |
| pHPSC208 | p41894-ACGC-ATG-3xFLAG-PGK1*-5xCGG-YFP | 1 | This work, exp60 |
| pHPSC513 | pSB2273-HEL2 | 5 | This work, exp153 |
| pHPSC514 | pSB2273-HEL2-c64a | 5 | This work, exp153 |
| pHPSC515 | pSB2273-HEL2-delring | 5 | This work, exp153 |
| pHPSC516 | pSB2273-ASC1 | 5 | This work, exp153 |
| pHPSC517 | pSB2273-ASC1-h16 | 5 | This work, exp153 |
| pHPSC518 | pSB2273-ASC1-r38 | 5 | This work, exp153 |
| pHPSC519 | pSB2273-ASC1-w85 | 5 | This work, exp153 |
| pHPSC512 | Barcoded plasmid pool (see plasmid cloning methods) | 2,5 | This work, exp152 |

**Table S2:**
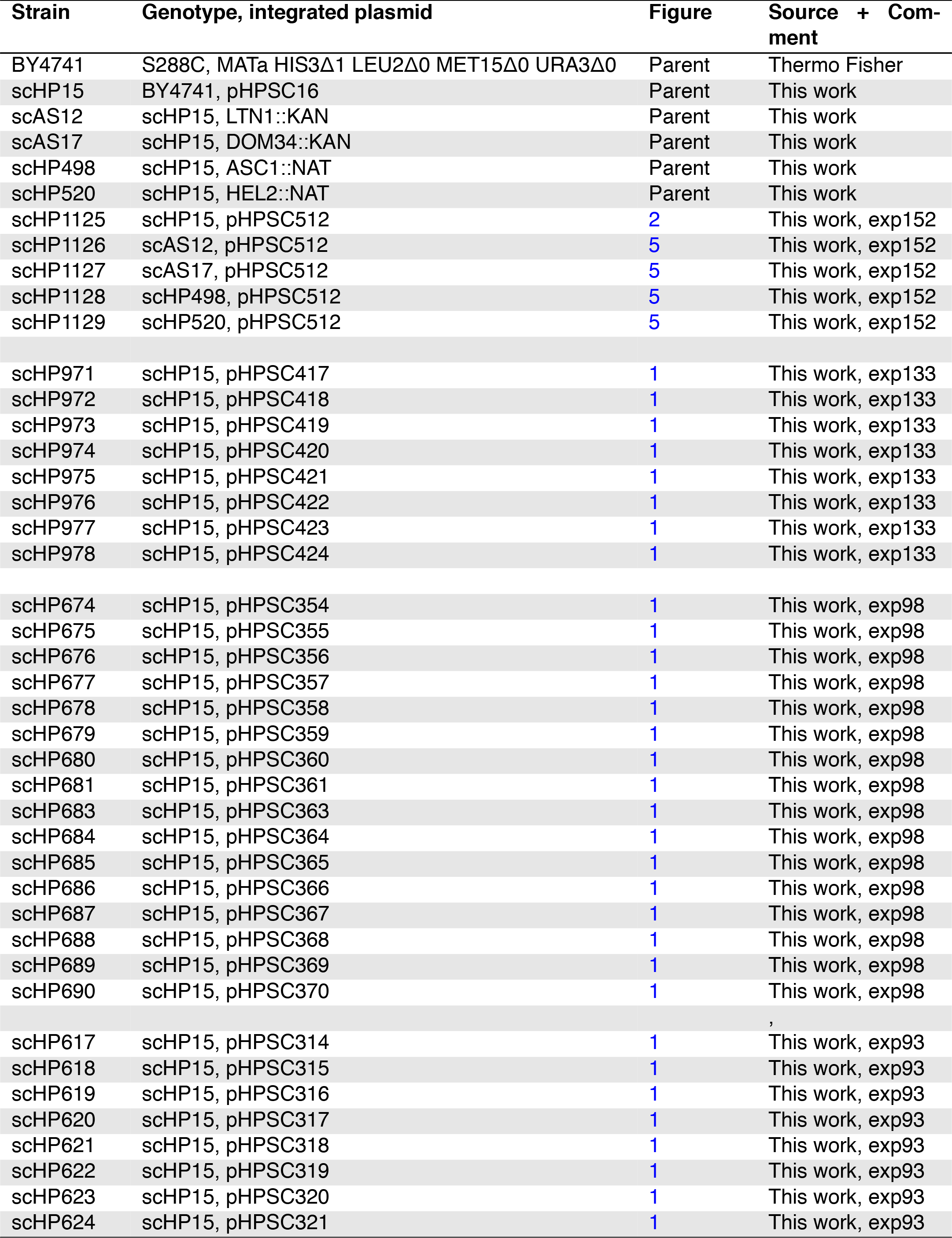
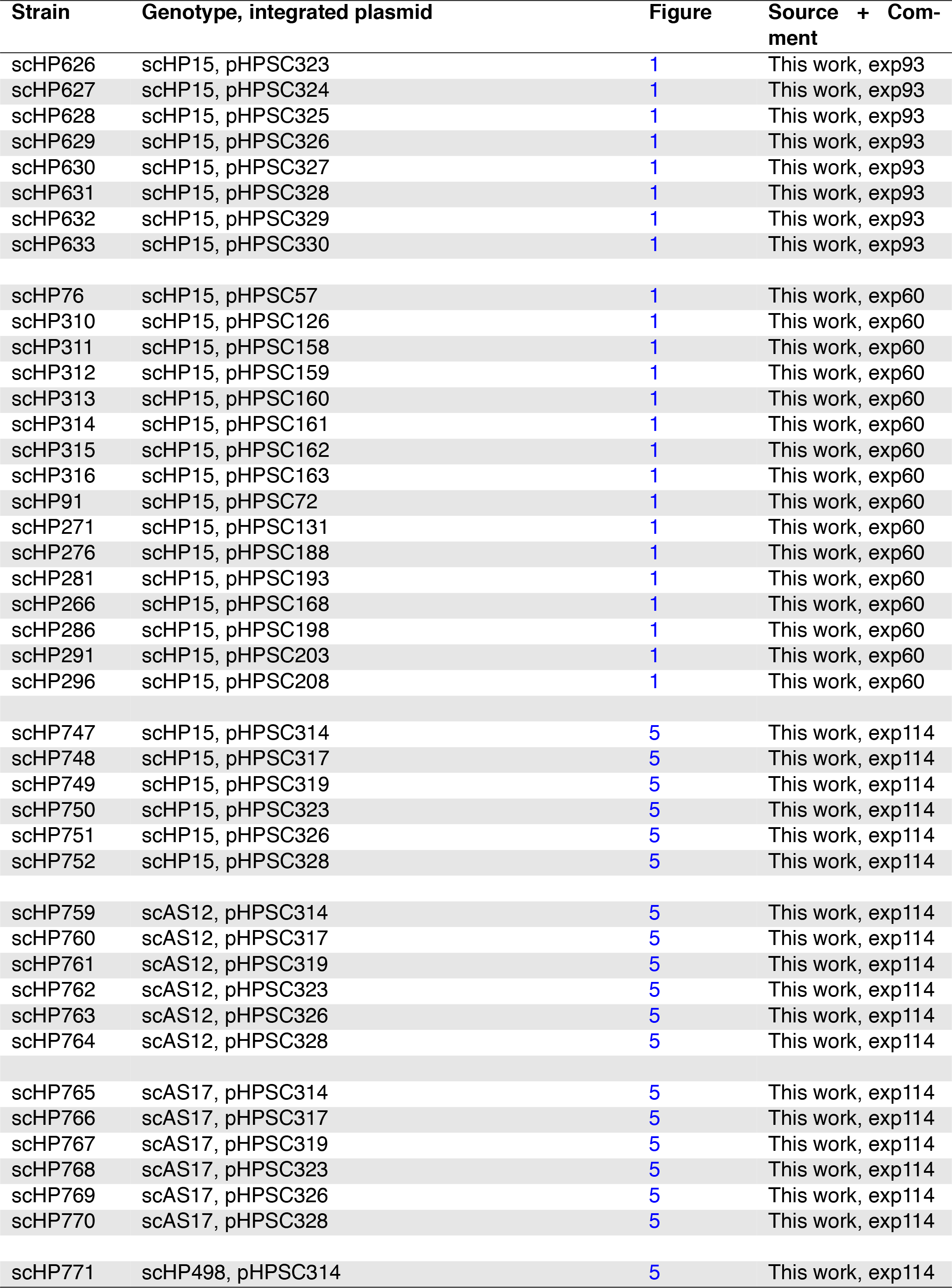
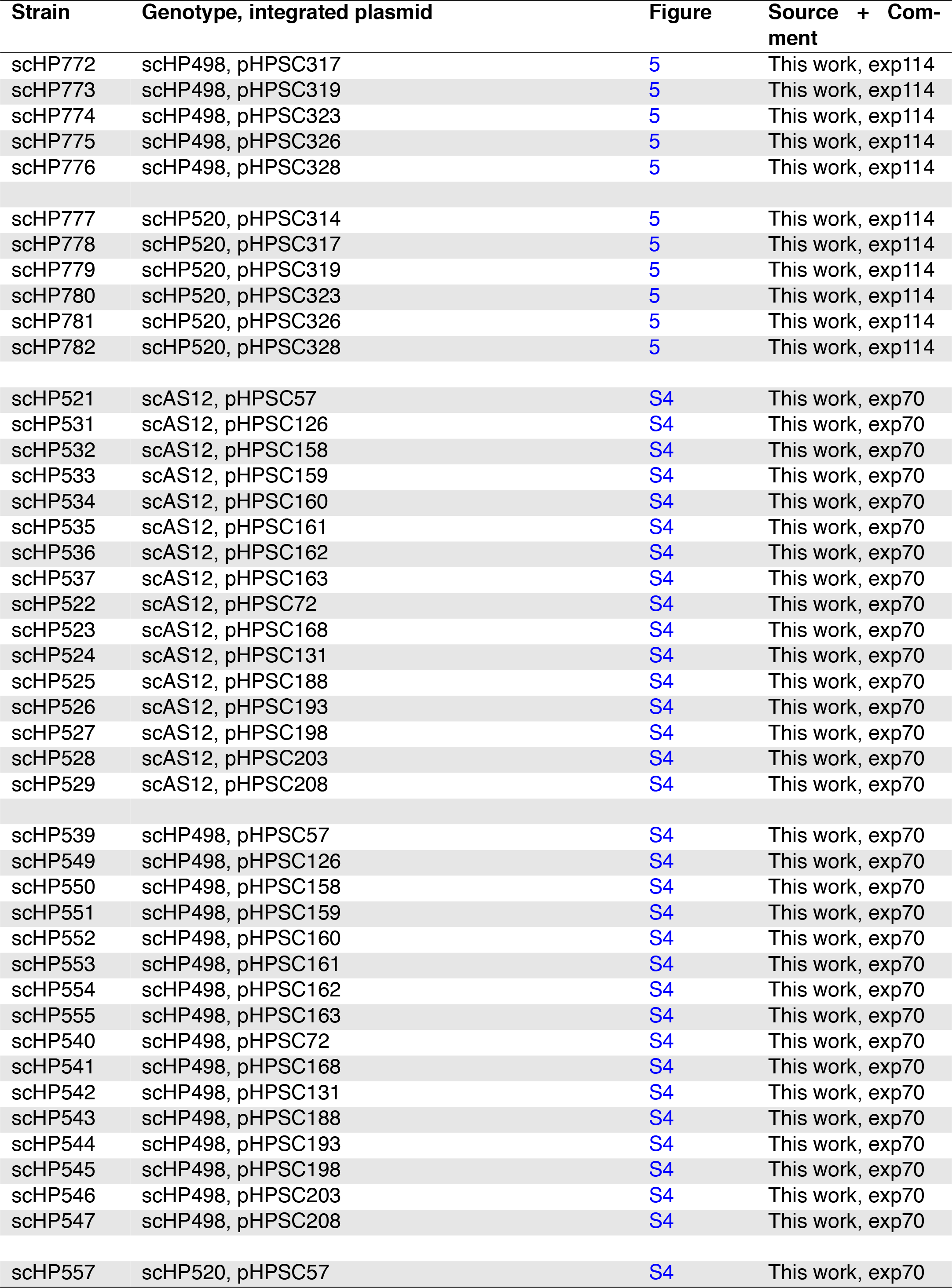
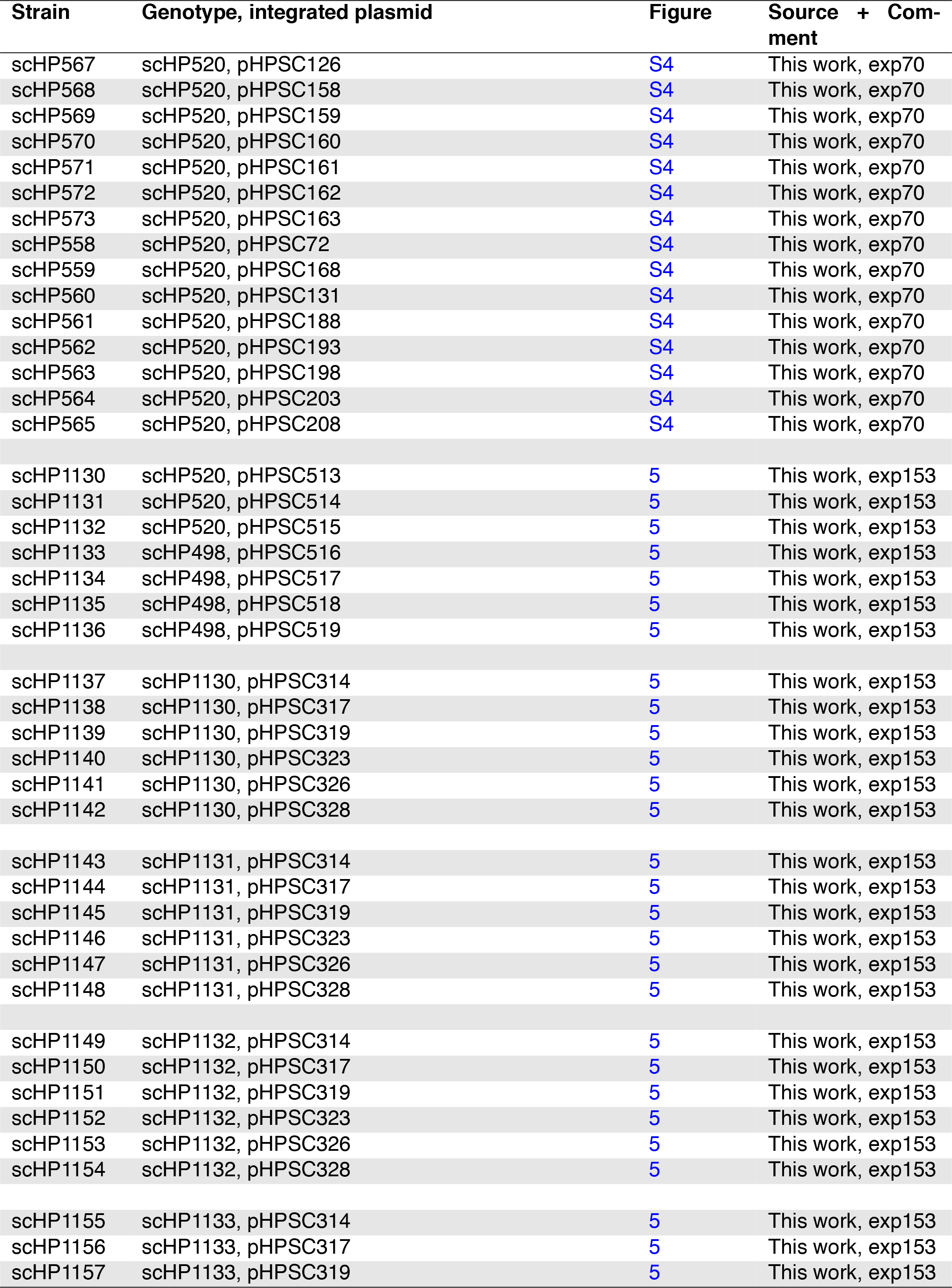
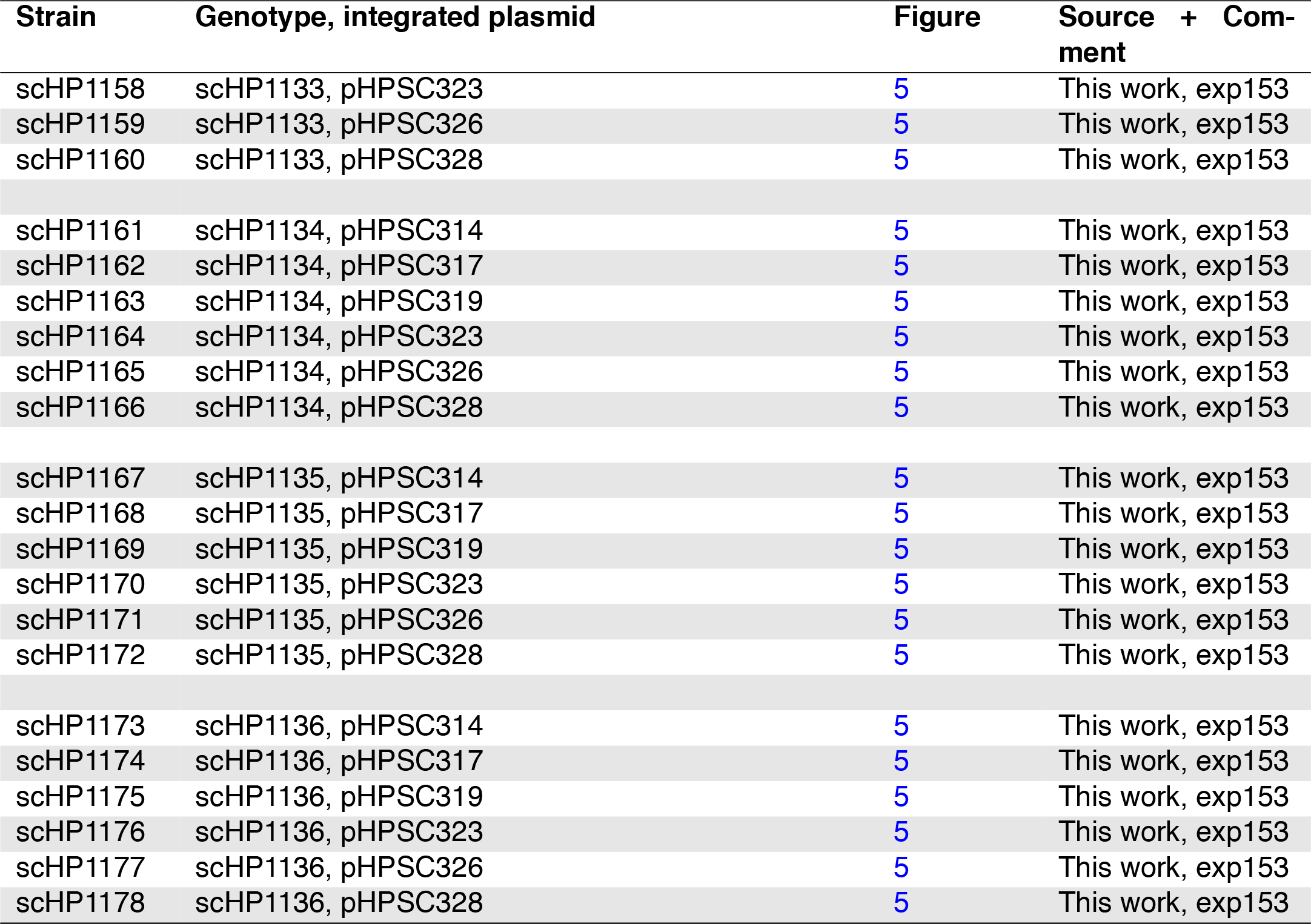
List of *S. cerevisiae* strains used in this study

| Strain | Genotype, integrated plasmid | Figure | Source + Comment |
| --- | --- | --- | --- |
| BY4741 | S288C, MATa HIS3Δ1 LEU2Δ0 MET15Δ0 URA3Δ0 | Parent | Thermo Fisher |
| scHP15 | BY4741, pHPSC16 | Parent | This work |
| scAS12 | scHP15, LTN1::KAN | Parent | This work |
| scAS17 | scHP15, DOM34::KAN | Parent | This work |
| scHP498 | scHP15, ASC1::NAT | Parent | This work |
| scHP520 | scHP15, HEL2::NAT | Parent | This work |
| scHP1125 | scHP15, pHPSC512 | 2 | This work, exp152 |
| scHP1126 | scAS12, pHPSC512 | 5 | This work, exp152 |
| scHP1127 | scAS17, pHPSC512 | 5 | This work, exp152 |
| scHP1128 | scHP498, pHPSC512 | 5 | This work, exp152 |
| scHP1129 | scHP520, pHPSC512 | 5 | This work, exp152 |
| scHP971 | scHP15, pHPSC417 | 1 | This work, exp133 |
| scHP972 | scHP15, pHPSC418 | 1 | This work, exp133 |
| scHP973 | scHP15, pHPSC419 | 1 | This work, exp133 |
| scHP974 | scHP15, pHPSC420 | 1 | This work, exp133 |
| scHP975 | scHP15, pHPSC421 | 1 | This work, exp133 |
| scHP976 | scHP15, pHPSC422 | 1 | This work, exp133 |
| scHP977 | scHP15, pHPSC423 | 1 | This work, exp133 |
| scHP978 | scHP15, pHPSC424 | 1 | This work, exp133 |
| scHP674 | scHP15, pHPSC354 | 1 | This work, exp98 |
| scHP675 | scHP15, pHPSC355 | 1 | This work, exp98 |
| scHP676 | scHP15, pHPSC356 | 1 | This work, exp98 |
| scHP677 | scHP15, pHPSC357 | 1 | This work, exp98 |
| scHP678 | scHP15, pHPSC358 | 1 | This work, exp98 |
| scHP679 | scHP15, pHPSC359 | 1 | This work, exp98 |
| scHP680 | scHP15, pHPSC360 | 1 | This work, exp98 |
| scHP681 | scHP15, pHPSC361 | 1 | This work, exp98 |
| scHP683 | scHP15, pHPSC363 | 1 | This work, exp98 |
| scHP684 | scHP15, pHPSC364 | 1 | This work, exp98 |
| scHP685 | scHP15, pHPSC365 | 1 | This work, exp98 |
| scHP686 | scHP15, pHPSC366 | 1 | This work, exp98 |
| scHP687 | scHP15, pHPSC367 | 1 | This work, exp98 |
| scHP688 | scHP15, pHPSC368 | 1 | This work, exp98 |
| scHP689 | scHP15, pHPSC369 | 1 | This work, exp98 |
| scHP690 | scHP15, pHPSC370 | 1 | This work, exp98 |
| scHP617 | scHP15, pHPSC314 | 1 | This work, exp93 |
| scHP618 | scHP15, pHPSC315 | 1 | This work, exp93 |
| scHP619 | scHP15, pHPSC316 | 1 | This work, exp93 |
| scHP620 | scHP15, pHPSC317 | 1 | This work, exp93 |
| scHP621 | scHP15, pHPSC318 | 1 | This work, exp93 |
| scHP622 | scHP15, pHPSC319 | 1 | This work, exp93 |
| scHP623 | scHP15, pHPSC320 | 1 | This work, exp93 |
| scHP624 | scHP15, pHPSC321 | 1 | This work, exp93 |
| scHP626 | scHP15, pHPSC323 | 1 | This work, exp93 |
| scHP627 | scHP15, pHPSC324 | 1 | This work, exp93 |
| scHP628 | scHP15, pHPSC325 | 1 | This work, exp93 |
| scHP629 | scHP15, pHPSC326 | 1 | This work, exp93 |
| scHP630 | scHP15, pHPSC327 | 1 | This work, exp93 |
| scHP631 | scHP15, pHPSC328 | 1 | This work, exp93 |
| scHP632 | scHP15, pHPSC329 | 1 | This work, exp93 |
| scHP633 | scHP15, pHPSC330 | 1 | This work, exp93 |
| scHP76 | scHP15, pHPSC57 | 1 | This work, exp60 |
| scHP310 | scHP15, pHPSC126 | 1 | This work, exp60 |
| scHP311 | scHP15, pHPSC158 | 1 | This work, exp60 |
| scHP312 | scHP15, pHPSC159 | 1 | This work, exp60 |
| scHP313 | scHP15, pHPSC160 | 1 | This work, exp60 |
| scHP314 | scHP15, pHPSC161 | 1 | This work, exp60 |
| scHP315 | scHP15, pHPSC162 | 1 | This work, exp60 |
| scHP316 | scHP15, pHPSC163 | 1 | This work, exp60 |
| scHP91 | scHP15, pHPSC72 | 1 | This work, exp60 |
| scHP271 | scHP15, pHPSC131 | 1 | This work, exp60 |
| scHP276 | scHP15, pHPSC188 | 1 | This work, exp60 |
| scHP281 | scHP15, pHPSC193 | 1 | This work, exp60 |
| scHP266 | scHP15, pHPSC168 | 1 | This work, exp60 |
| scHP286 | scHP15, pHPSC198 | 1 | This work, exp60 |
| scHP291 | scHP15, pHPSC203 | 1 | This work, exp60 |
| scHP296 | scHP15, pHPSC208 | 1 | This work, exp60 |
| scHP747 | scHP15, pHPSC314 | 5 | This work, exp114 |
| scHP748 | scHP15, pHPSC317 | 5 | This work, exp114 |
| scHP749 | scHP15, pHPSC319 | 5 | This work, exp114 |
| scHP750 | scHP15, pHPSC323 | 5 | This work, exp114 |
| scHP751 | scHP15, pHPSC326 | 5 | This work, exp114 |
| scHP752 | scHP15, pHPSC328 | 5 | This work, exp114 |
| scHP759 | scAS12, pHPSC314 | 5 | This work, exp114 |
| scHP760 | scAS12, pHPSC317 | 5 | This work, exp114 |
| scHP761 | scAS12, pHPSC319 | 5 | This work, exp114 |
| scHP762 | scAS12, pHPSC323 | 5 | This work, exp114 |
| scHP763 | scAS12, pHPSC326 | 5 | This work, exp114 |
| scHP764 | scAS12, pHPSC328 | 5 | This work, exp114 |
| scHP765 | scAS17, pHPSC314 | 5 | This work, exp114 |
| scHP766 | scAS17, pHPSC317 | 5 | This work, exp114 |
| scHP767 | scAS17, pHPSC319 | 5 | This work, exp114 |
| scHP768 | scAS17, pHPSC323 | 5 | This work, exp114 |
| scHP769 | scAS17, pHPSC326 | 5 | This work, exp114 |
| scHP770 | scAS17, pHPSC328 | 5 | This work, exp114 |
| scHP771 | scHP498, pHPSC314 | 5 | This work, exp114 |

| Continued from previous page |  |  |  |
| --- | --- | --- | --- |
| Strain | Genotype, integrated plasmid | Figure | Source + Comment |
| scHP772 | scHP498, pHPSC317 | 5 | This work, exp114 |
| scHP773 | scHP498, pHPSC319 | 5 | This work, exp114 |
| scHP774 | scHP498, pHPSC323 | 5 | This work, exp114 |
| scHP775 | scHP498, pHPSC326 | 5 | This work, exp114 |
| scHP776 | scHP498, pHPSC328 | 5 | This work, exp114 |
| scHP777 | scHP520, pHPSC314 | 5 | This work, exp114 |
| scHP778 | scHP520, pHPSC317 | 5 | This work, exp114 |
| scHP779 | scHP520, pHPSC319 | 5 | This work, exp114 |
| scHP780 | scHP520, pHPSC323 | 5 | This work, exp114 |
| scHP781 | scHP520, pHPSC326 | 5 | This work, exp114 |
| scHP782 | scHP520, pHPSC328 | 5 | This work, exp114 |
| scHP521 | scAS12, pHPSC57 | S4 | This work, exp70 |
| scHP531 | scAS12, pHPSC126 | S4 | This work, exp70 |
| scHP532 | scAS12, pHPSC158 | S4 | This work, exp70 |
| scHP533 | scAS12, pHPSC159 | S4 | This work, exp70 |
| scHP534 | scAS12, pHPSC160 | S4 | This work, exp70 |
| scHP535 | scAS12, pHPSC161 | S4 | This work, exp70 |
| scHP536 | scAS12, pHPSC162 | S4 | This work, exp70 |
| scHP537 | scAS12, pHPSC163 | S4 | This work, exp70 |
| scHP522 | scAS12, pHPSC72 | S4 | This work, exp70 |
| scHP523 | scAS12, pHPSC168 | S4 | This work, exp70 |
| scHP524 | scAS12, pHPSC131 | S4 | This work, exp70 |
| scHP525 | scAS12, pHPSC188 | S4 | This work, exp70 |
| scHP526 | scAS12, pHPSC193 | S4 | This work, exp70 |
| scHP527 | scAS12, pHPSC198 | S4 | This work, exp70 |
| scHP528 | scAS12, pHPSC203 | S4 | This work, exp70 |
| scHP529 | scAS12, pHPSC208 | S4 | This work, exp70 |
| scHP539 | scHP498, pHPSC57 | S4 | This work, exp70 |
| scHP549 | scHP498, pHPSC126 | S4 | This work, exp70 |
| scHP550 | scHP498, pHPSC158 | S4 | This work, exp70 |
| scHP551 | scHP498, pHPSC159 | S4 | This work, exp70 |
| scHP552 | scHP498, pHPSC160 | S4 | This work, exp70 |
| scHP553 | scHP498, pHPSC161 | S4 | This work, exp70 |
| scHP554 | scHP498, pHPSC162 | S4 | This work, exp70 |
| scHP555 | scHP498, pHPSC163 | S4 | This work, exp70 |
| scHP540 | scHP498, pHPSC72 | S4 | This work, exp70 |
| scHP541 | scHP498, pHPSC168 | S4 | This work, exp70 |
| scHP542 | scHP498, pHPSC131 | S4 | This work, exp70 |
| scHP543 | scHP498, pHPSC188 | S4 | This work, exp70 |
| scHP544 | scHP498, pHPSC193 | S4 | This work, exp70 |
| scHP545 | scHP498, pHPSC198 | S4 | This work, exp70 |
| scHP546 | scHP498, pHPSC203 | S4 | This work, exp70 |
| scHP547 | scHP498, pHPSC208 | S4 | This work, exp70 |
| scHP557 | scHP520, pHPSC57 | S4 | This work, exp70 |
| scHP567 | scHP520, pHPSC126 | S4 | This work, exp70 |
| scHP568 | scHP520, pHPSC158 | S4 | This work, exp70 |
| scHP569 | scHP520, pHPSC159 | S4 | This work, exp70 |
| scHP570 | scHP520, pHPSC160 | S4 | This work, exp70 |
| scHP571 | scHP520, pHPSC161 | S4 | This work, exp70 |
| scHP572 | scHP520, pHPSC162 | S4 | This work, exp70 |
| scHP573 | scHP520, pHPSC163 | S4 | This work, exp70 |
| scHP558 | scHP520, pHPSC72 | S4 | This work, exp70 |
| scHP559 | scHP520, pHPSC168 | S4 | This work, exp70 |
| scHP560 | scHP520, pHPSC131 | S4 | This work, exp70 |
| scHP561 | scHP520, pHPSC188 | S4 | This work, exp70 |
| scHP562 | scHP520, pHPSC193 | S4 | This work, exp70 |
| scHP563 | scHP520, pHPSC198 | S4 | This work, exp70 |
| scHP564 | scHP520, pHPSC203 | S4 | This work, exp70 |
| scHP565 | scHP520, pHPSC208 | S4 | This work, exp70 |
| scHP1130 | scHP520, pHPSC513 | 5 | This work, exp153 |
| scHP1131 | scHP520, pHPSC514 | 5 | This work, exp153 |
| scHP1132 | scHP520, pHPSC515 | 5 | This work, exp153 |
| scHP1133 | scHP498, pHPSC516 | 5 | This work, exp153 |
| scHP1134 | scHP498, pHPSC517 | 5 | This work, exp153 |
| scHP1135 | scHP498, pHPSC518 | 5 | This work, exp153 |
| scHP1136 | scHP498, pHPSC519 | 5 | This work, exp153 |
| scHP1137 | scHP1130, pHPSC314 | 5 | This work, exp153 |
| scHP1138 | scHP1130, pHPSC317 | 5 | This work, exp153 |
| scHP1139 | scHP1130, pHPSC319 | 5 | This work, exp153 |
| scHP1140 | scHP1130, pHPSC323 | 5 | This work, exp153 |
| scHP1141 | scHP1130, pHPSC326 | 5 | This work, exp153 |
| scHP1142 | scHP1130, pHPSC328 | 5 | This work, exp153 |
| scHP1143 | scHP1131, pHPSC314 | 5 | This work, exp153 |
| scHP1144 | scHP1131, pHPSC317 | 5 | This work, exp153 |
| scHP1145 | scHP1131, pHPSC319 | 5 | This work, exp153 |
| scHP1146 | scHP1131, pHPSC323 | 5 | This work, exp153 |
| scHP1147 | scHP1131, pHPSC326 | 5 | This work, exp153 |
| scHP1148 | scHP1131, pHPSC328 | 5 | This work, exp153 |
| scHP1149 | scHP1132, pHPSC314 | 5 | This work, exp153 |
| scHP1150 | scHP1132, pHPSC317 | 5 | This work, exp153 |
| scHP1151 | scHP1132, pHPSC319 | 5 | This work, exp153 |
| scHP1152 | scHP1132, pHPSC323 | 5 | This work, exp153 |
| scHP1153 | scHP1132, pHPSC326 | 5 | This work, exp153 |
| scHP1154 | scHP1132, pHPSC328 | 5 | This work, exp153 |
| scHP1155 | scHP1133, pHPSC314 | 5 | This work, exp153 |
| scHP1156 | scHP1133, pHPSC317 | 5 | This work, exp153 |
| scHP1157 | scHP1133, pHPSC319 | 5 | This work, exp153 |

| Continued from previous page |  |  |  |
| --- | --- | --- | --- |
| Strain | Genotype, integrated plasmid | Figure | Source + Comment |
| scHP1158 | scHP1133, pHPSC323 | 5 | This work, exp153 |
| scHP1159 | scHP1133, pHPSC326 | 5 | This work, exp153 |
| scHP1160 | scHP1133, pHPSC328 | 5 | This work, exp153 |
| scHP1161 | scHP1134, pHPSC314 | 5 | This work, exp153 |
| scHP1162 | scHP1134, pHPSC317 | 5 | This work, exp153 |
| scHP1163 | scHP1134, pHPSC319 | 5 | This work, exp153 |
| scHP1164 | scHP1134, pHPSC323 | 5 | This work, exp153 |
| scHP1165 | scHP1134, pHPSC326 | 5 | This work, exp153 |
| scHP1166 | scHP1134, pHPSC328 | 5 | This work, exp153 |
| scHP1167 | scHP1135, pHPSC314 | 5 | This work, exp153 |
| scHP1168 | scHP1135, pHPSC317 | 5 | This work, exp153 |
| scHP1169 | scHP1135, pHPSC319 | 5 | This work, exp153 |
| scHP1170 | scHP1135, pHPSC323 | 5 | This work, exp153 |
| scHP1171 | scHP1135, pHPSC326 | 5 | This work, exp153 |
| scHP1172 | scHP1135, pHPSC328 | 5 | This work, exp153 |
| scHP1173 | scHP1136, pHPSC314 | 5 | This work, exp153 |
| scHP1174 | scHP1136, pHPSC317 | 5 | This work, exp153 |
| scHP1175 | scHP1136, pHPSC319 | 5 | This work, exp153 |
| scHP1176 | scHP1136, pHPSC323 | 5 | This work, exp153 |
| scHP1177 | scHP1136, pHPSC326 | 5 | This work, exp153 |
| scHP1178 | scHP1136, pHPSC328 | 5 | This work, exp153 |

**Table S3:**
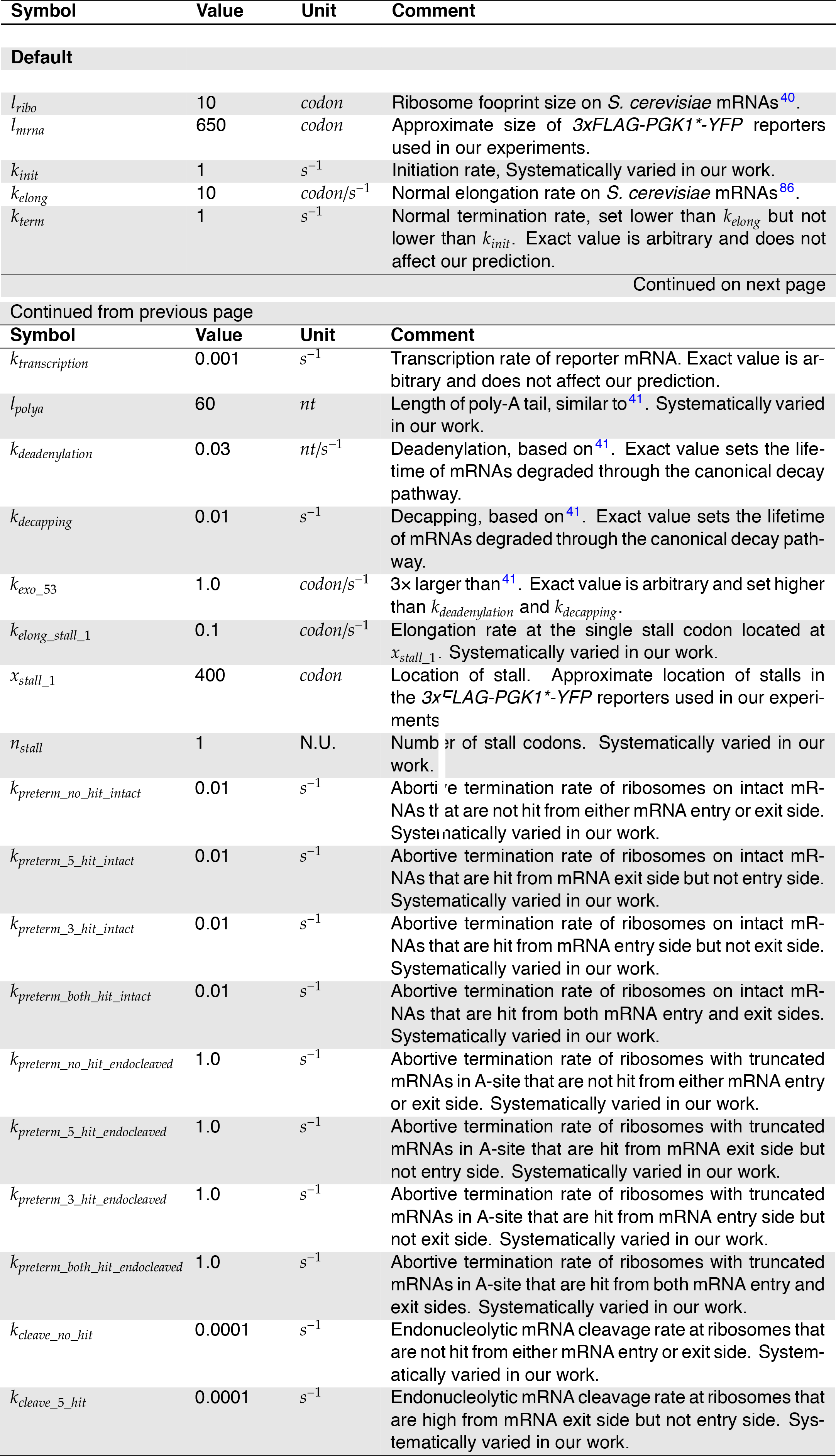
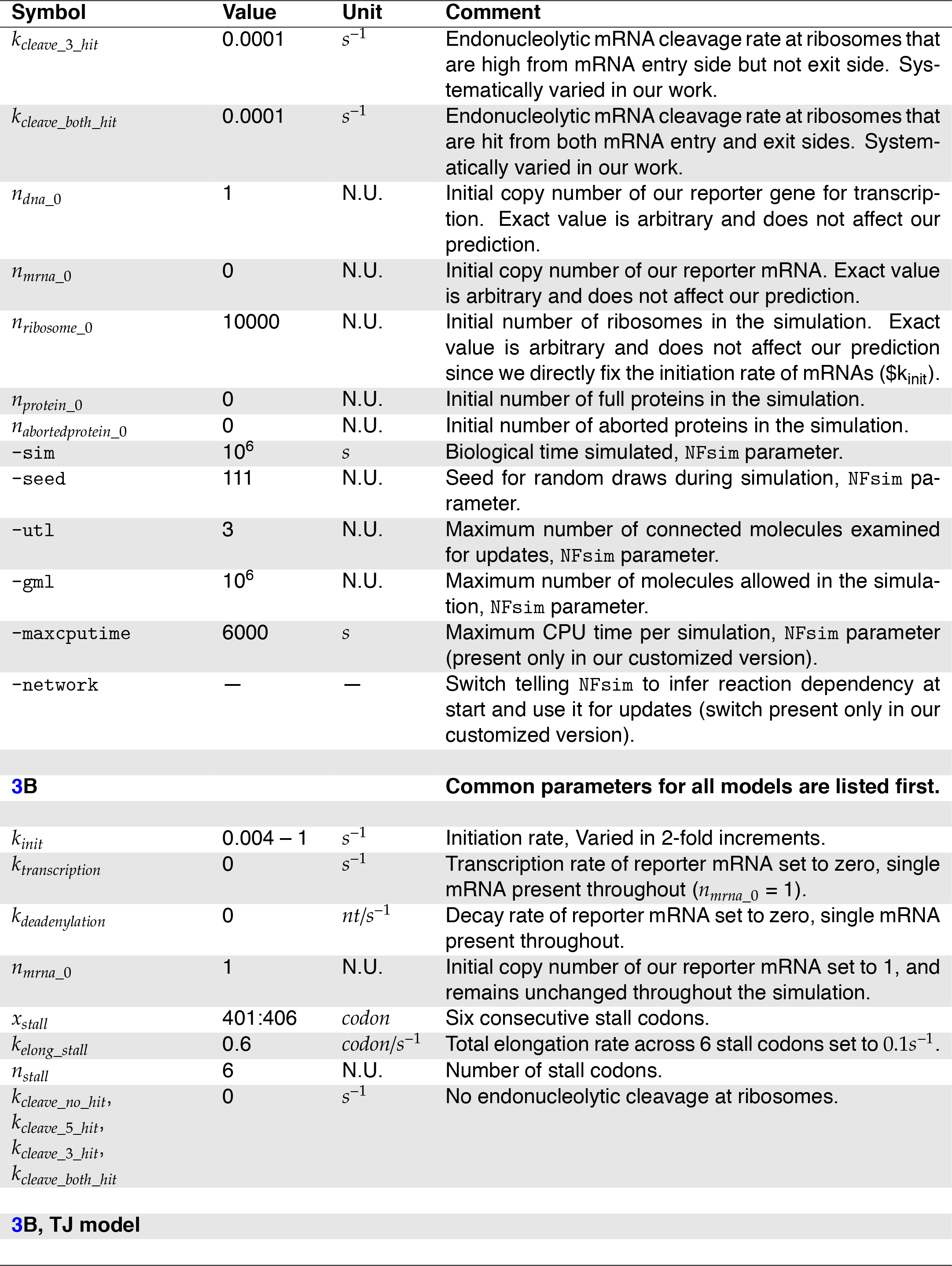
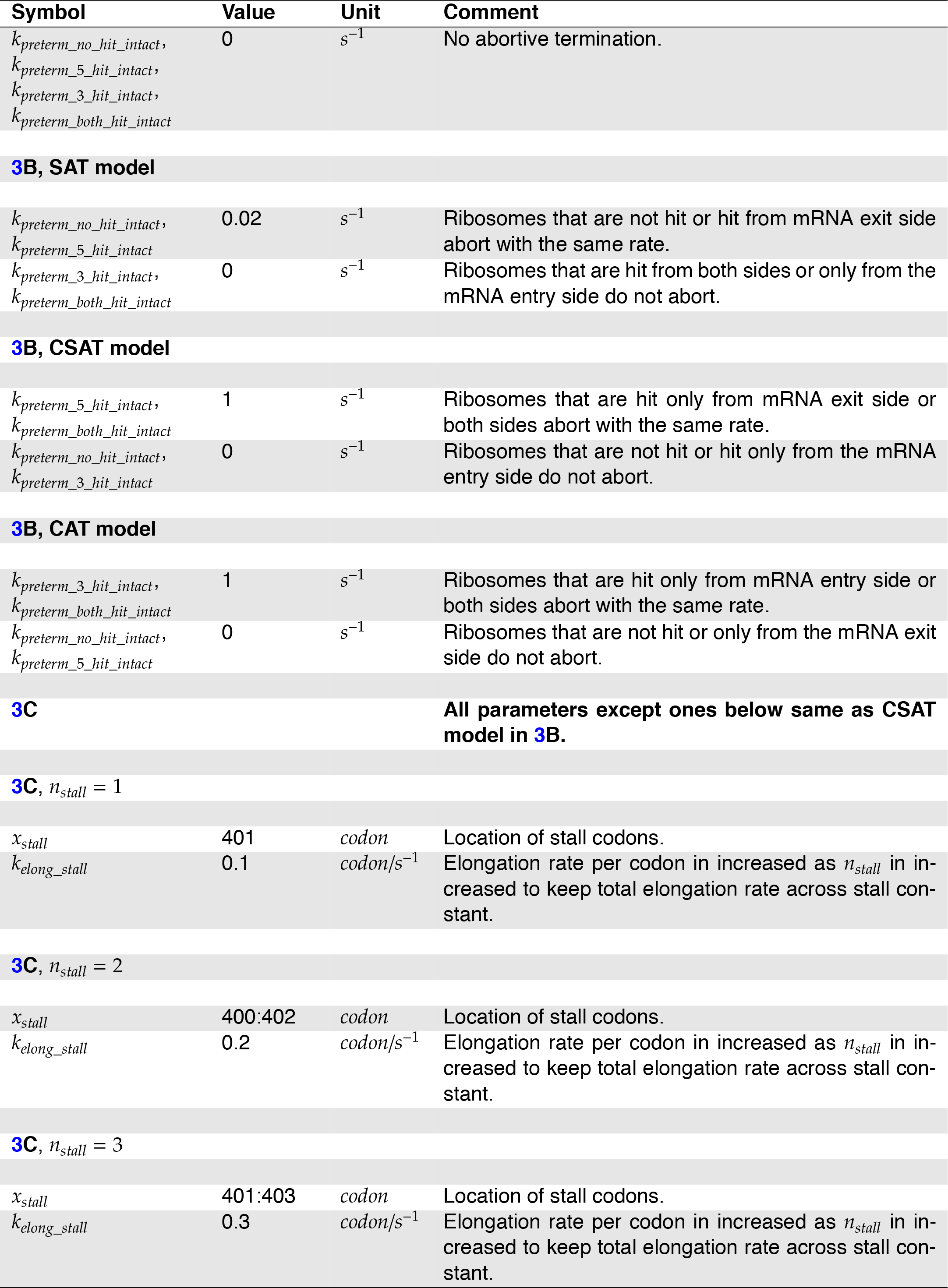
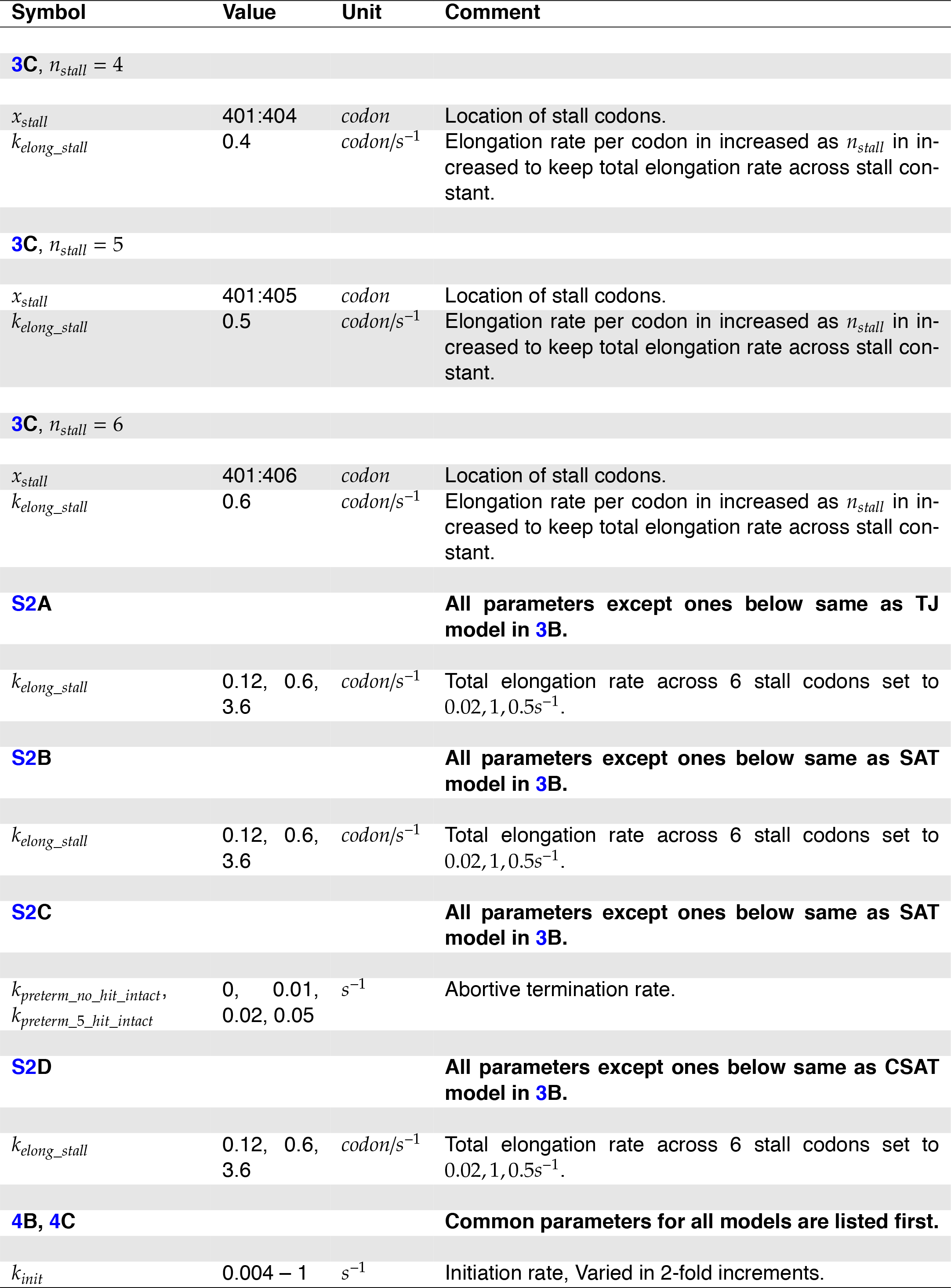
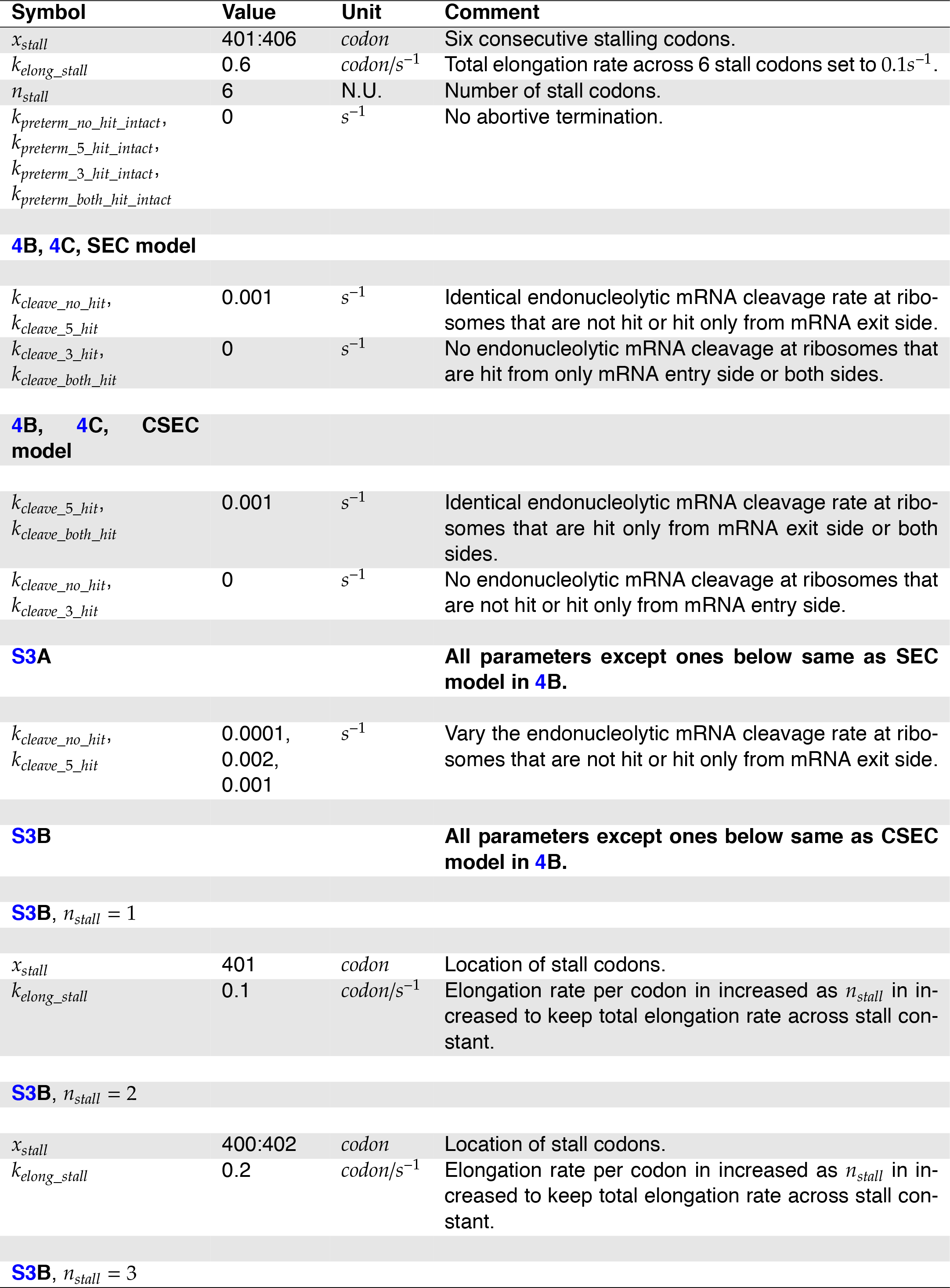
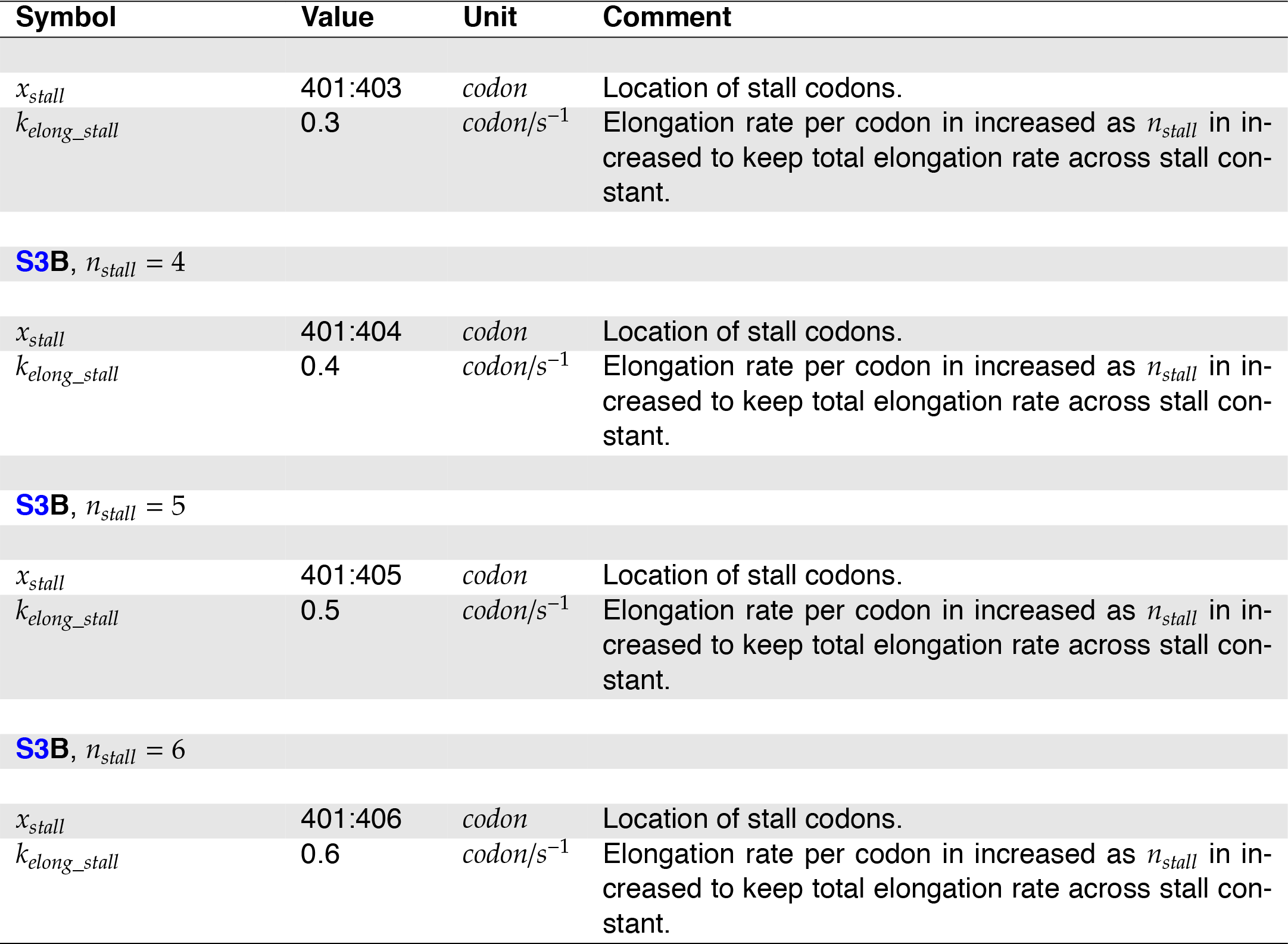
Simulation Parameters. Default parameter values across all simulations are listed first. Parameters altered for specific simulations are listed next under the corresponding figure. Parameters not listed for a specific Figure are either set to the default value or explicitly indicated to be the same as for another figure. N.U. stands for No Units. The symbol names correspond to the ones used in `tasep.py`. For parameters that are not systematically varied in our work but that have been previously measured, we chose them to be a convenient rounded value within 2-fold of the measured value. For example, we set the ribosome footprint size to 10 codons while it is closer to 9 codons ^40^; The decapping rate of *PGK1* mRNA is set to 0.01*s*^−1^ while it is measured to be 0.008*s* ^−1 41^. We do not expect these choices to alter any of the results presented in this study.

